# CTP regulates membrane-binding activity of the nucleoid occlusion protein Noc

**DOI:** 10.1101/2021.02.11.430593

**Authors:** Adam S. B. Jalal, Ngat T. Tran, Ling J. Wu, Karunakaran Ramakrishnan, Martin Rejzek, Giulia Gobbato, Clare E. M. Stevenson, David M. Lawson, Jeff Errington, Tung B. K. Le

**Affiliations:** Department of Molecular Microbiology John Innes Centre, Norwich, NR4 7UH, United Kingdom; Centre for Bacterial Cell Biology, Biosciences Institute, Medical School Newcastle University, Newcastle Upon Tyne, NE2 4AX, United Kingdom; Chemistry Platform John Innes Centre, Norwich, NR4 7UH, United Kingdom; Department of Biological Chemistry John Innes Centre, Norwich, NR4 7UH, United Kingdom

## Abstract

ATP and GTP-dependent molecular switches are extensively used to control functions of proteins in a wide range of biological processes. However, CTP switches are rarely reported. Here, we report that a nucleoid occlusion protein Noc is a CTPase enzyme whose membrane-binding activity is directly regulated by a CTP switch. In *Bacillus subtilis*, Noc nucleates on 16-bp *NBS* sites before associating with neighboring non-specific DNA to form large membrane-associated nucleoprotein complexes to physically occlude assembly of the cell division machinery. By *in vitro* reconstitution, we show that (i) CTP is required for Noc to form the *NBS*-dependent nucleoprotein complex, and (ii) CTP binding, but not hydrolysis, switches Noc to a membrane-active state. Overall, we suggest that CTP couples membrane-binding activity of Noc to nucleoprotein complex formation to ensure productive recruitment of DNA to the bacterial cell membrane for nucleoid occlusion activity.

## INTRODUCTION

While ATP and GTP switches are ubiquitous in biology, CTP switches have rarely been identified but may be more widespread than previously appreciated. A recent discovery showed that ParB, a crucial protein for bacterial chromosome segregation, is the founding member of a new class of CTP-dependent molecular switches ^1,2^. ParB nucleates on a *parS* DNA sequence and associates with neighboring DNA, a process known as spreading, to enable faithful chromosome segregation ^3–8^. CTP induces ParB self-dimerization to create a clamp-like molecule ^2^. The ParB clamp self-loads at *parS*, then spreads by sliding to neighboring DNA while still entrapping DNA ^2,9^. Essentially, CTP serves to switch ParB from a *parS*-nucleating open clamp to a DNA-sliding closed clamp state ^2,9^. The result is the formation of a higher-order nucleoprotein complex with multiple ParB-CTP clamps entrapped in the vicinity of the *parS* locus. The higher-order nucleoprotein complex stimulates the ATPase activity of ParA, a partner of ParB, driving the segregation of replicated chromosomes to daughter cells ^3,10–14^.

In Firmicutes, the nucleoid occlusion protein Noc is a paralog of ParB ^15–17^; however, the role of Noc is different from that of a canonical ParB ^18–20^. Noc helps to direct the assembly of the cell division machinery towards the middle of a dividing cell where the concentration of chromosomal DNA (the nucleoid) is the least, thus ensuring a binary cell division ^18,21–23^. Noc does so by nucleating on 16-bp *NBS* (Noc-binding site) sites scattered around the chromosome before spreading to neighboring DNA to form large Noc-DNA nucleoprotein complexes ^24^. Unusually, Noc is also a peripheral membrane protein that directly associates with the cell membrane via a predicted N-terminal amphipathic helix ^21^ (Figure 1A). The recruitment of the chromosomal DNA to the membrane is crucial for preventing the assembly of the division machinery over the chromosome; indeed a Noc variant lacking the amphipathic helix is impaired in nucleoid occlusion activity ^21^. In *Bacillus subtilis*, Noc was observed to associate with the cell membrane in a transient manner *in vivo* ^24^. It was thought that a strong membrane-binding activity of Noc might have been selected against since a stable association with the membrane might hamper chromosome replication and segregation ^21^, yet it is unclear how the membrane-binding activity of Noc is modulated. Furthermore, Noc must bring the chromosomal DNA to the membrane to physically inhibit the assembly of the division machinery ^21^; an unregulated membrane-binding activity would likely confine apo-Noc permanently to the cell membrane, thus unfavorably limiting the recruitment of DNA to the membrane ^21^. Again, it remains unclear whether the membrane-binding activity of Noc is regulated, and if so how.

**Figure 1.**
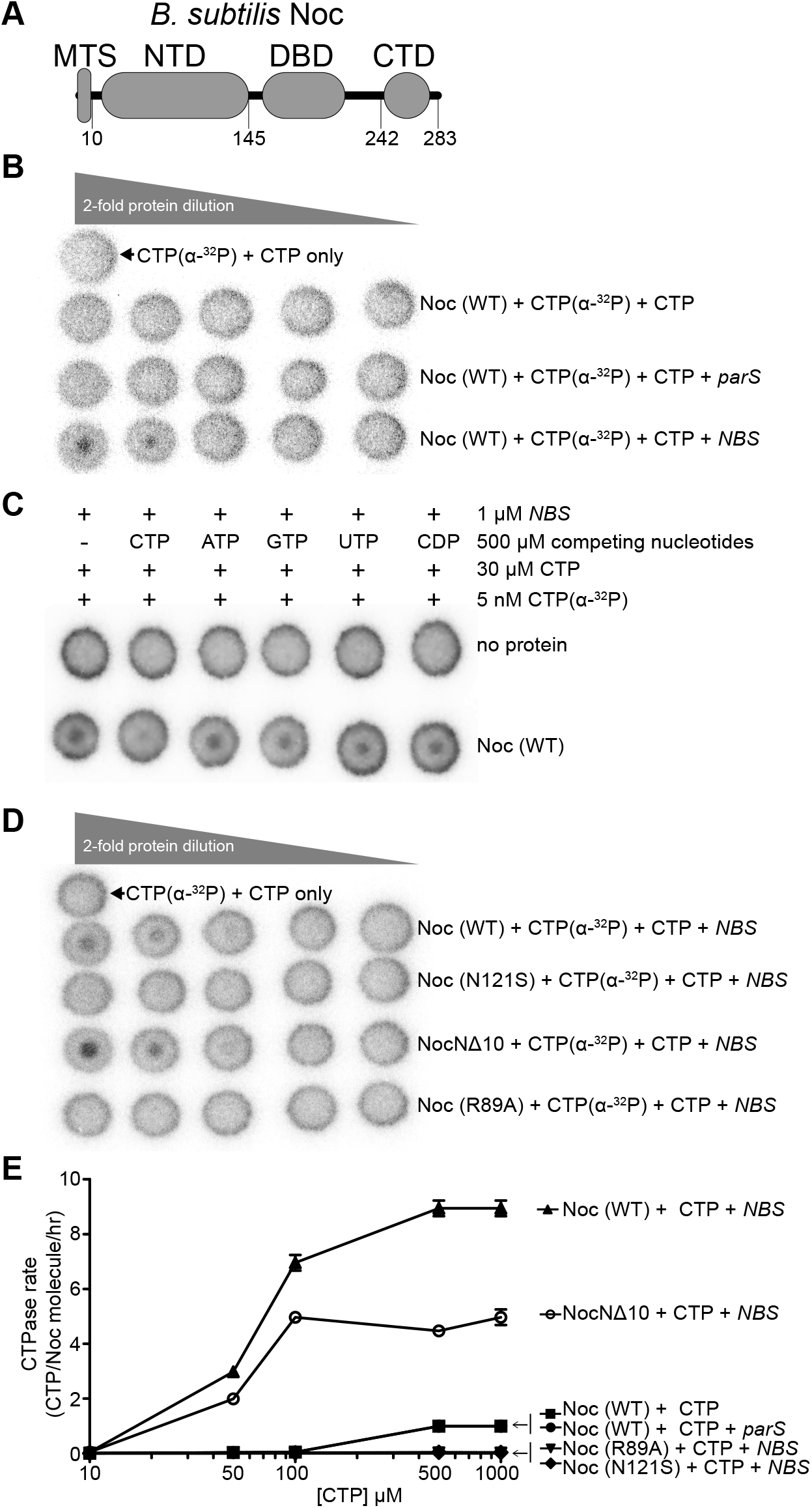
Noc binds and hydrolyzes CTP in the presence of *NBS* DNA. **(A)** The domain architecture of *B. subtilis* Noc: a membrane-targeting sequence (MTS), an N-terminal domain (NTD), a central DNA-binding domain (DBD), and a C-terminal domain (CTD). **(B-D)** CTP binding as monitored by DRaCALA assay using radiolabeled CTP α-P^32^. The bulls-eye staining indicates CTP binding due to a more rapid immobilization of protein-ligand complexes compared to free ligands. The starting concentration of Noc used in all panels was 30 μM. The concentrations of CTP α-P^32^, unlabeled CTP, and a 22-bp *parS*/*NBS* DNA used in all panels were 5 nM, 30 μM, and 1 μM, respectively. **(E)** CTP hydrolysis rates of Noc (WT) and variants were measured by continuous detection of released inorganic phosphates (see STAR Methods). CTPase rates were measured at increasing concentrations of CTP. All reactions contained 1 μM Noc (WT/variants) ± 1 μM 22-bp *NBS* or *parS* DNA and an increasing concentration of CTP (0, 10, 50, 100, 500, and 1000 μM). Experiments were triplicated and the standard deviation of the CTPase rates were presented.

To investigate further, we have biochemically reconstituted *NBS*-dependent Noc spreading and membrane association events using purified *B. subtilis* Noc protein and phospholipid vesicles. We show that, similar to a canonical ParB, Noc is a CTPase enzyme that binds CTP to form a protein clamp that can slide and entrap DNA. Importantly, CTP binding, but not hydrolysis, is required to switch Noc-DNA from a membrane inactive to an active state, thus locking Noc into a pathway in which it has to spread before associating with the cell membrane. We solve an X-ray crystal structure of a C-terminal domain truncated apo-Noc from *Geobacillus thermoleovorans*, in which its membrane-targeting amphipathic helix adopts an autoinhibitory conformation, restricted from interacting with the membrane. We suggest that CTP binding might liberate the amphipathic helix, thereby switching Noc to a membrane-active state. Altogether, we demonstrate that CTP directly regulates the membrane-binding activity of the nucleoid occlusion protein Noc, further expanding the role of CTP switches in biology.

## RESULTS

### *NBS* DNA increases the CTP binding and hydrolysis rate of Noc

Given the shared ancestry between ParB and Noc, we wondered if *B. subtilis* Noc also binds and hydrolyses CTP. To investigate, we employed a membrane-spotting assay (DRaCALA) and the result showed that *B. subtilis* Noc binds radiolabeled CTP but only in the presence of a cognate 22-bp *NBS* DNA (Figure 1B). An excess of unlabeled CTP, but no other NTP or CDP, outcompeted radiolabeled CTP for binding to Noc, suggesting that *B. subtilis* Noc binds CTP specifically (Figure 1C). Similarly, an N-terminally truncated Noc variant lacking the 10-AA membrane targeting sequence (NocNΔ10) also bound radiolabeled CTP in the presence of *NBS* DNA (Figure 1D). However, the Noc (R89A) and Noc (N121S) variants, whose equivalent substitutions in ParB have been shown to impair spreading and CTP binding ^1,2,9^ (Figure S1A), did not bind radiolabeled CTP at the tested concentration (Figure 1D). Next, we performed a quantitative nucleotide-binding assay using isothermal titration calorimetry (ITC) with a non-hydrolyzable CTP analog (CTPγS) to ensure the heat exchange was solely due to nucleotide-binding but not hydrolysis or *NBS* DNA binding. We found that *B. subtilis* Noc binds CTPγS with a moderate affinity (K_d_ = 68 ± 23 μM), while Noc (N121S) bound CTPγS weaker at K_d_ = 232 ± 66 μM, and Noc (R89A) did not detectably bind nucleotide (Figure S1B). Consistent with a previous report ^2^, *B. subtilis* Noc also showed CTP hydrolysis activity, albeit at a low rate of ~1 CTP per Noc per hour when only the purified protein and CTP were included (Figure 1E) ^2^. The addition of a 22-bp *NBS* DNA, but not a non-cognate 22-bp *parS* DNA, increased the CTP hydrolysis rate ninefold to ~9 CTP per Noc per hour (Figure 1E) ^2^. The Noc (R89A) and Noc (N121S) variants did not show noticeable CTP-hydrolyzing activity (Figure 1E). Lastly, despite binding CTP equally or more strongly than the WT (Figure 1D), NocNΔ10 hydrolyzed CTP at a reduced rate of ~5 CTP per Noc per hour (Figure 1E) (see Discussion). Altogether, our data suggest that *B. subtilis* Noc is a CTPase enzyme that binds and hydrolyses CTP in the presence of cognate DNA.

### CTP and *NBS* DNA stimulate the engagement of the N-terminal domain of Noc *in vitro*

In the presence of CTP, ParB self-engages at the N-terminal domain (NTD) to create a clamp-like molecule ^2,9^. To investigate whether CTP elicits a similar response in Noc, we employed site-specific crosslinking of a purified *B. subtilis* Noc (E29C) variant by a sulfhydryl-to-sulfhydryl crosslinker bismaleimidoethane (BMOE). Based on a sequence alignment between *B. subtilis* ParB and Noc (Figure S1A), residue E29 at the NTD was selected and substituted by cysteine on an otherwise cysteine-less Noc (WT) background (Figure S2A), to create a variant in which symmetry-related cysteines become covalently linked together if they are within 8 Å of each other. The crosslinked form was detectable as a protein form with reduced mobility on SDS-PAGE (labeled X in Figure 2). In the absence of CTP, Noc (E29C) crosslinked minimally (~10% crosslinked fraction, lanes 2-4 of Figure 2A). The crosslinking efficiency increased threefold (~30%) in the presence of CTP (lane 8 of Figure 2A), but not CDP or ATP (Figure 2A and Figure S2B). The crosslinking efficiency further increased fivefold (~55%) when both CTP and a 22-bp *NBS* were included (lane 10 of Figure 2A, and Figure S2D). However, the addition of a non-cognate 22-bp *parS* DNA did not result in the same high level of crosslinking, even when CTP was present (lane 9 of Figure 2A). Noticeably, non-hydrolyzable CTPγS readily promoted crosslinking (~45% crosslinked fraction) regardless of the presence or absence of *NBS* DNA (lanes 8-10 of Figure 2B, and Figure S2E). Therefore, our data suggest that CTP binding, but not hydrolysis, is required for the NTD engagement of *B. subtilis* Noc. Consistent with the requirement of CTP binding for NTD engagement, the Noc (E29C R89A) variant, in which the R89A substitution incapacitates CTP binding, did not crosslink beyond the background level in any tested condition (Figure 2C). The Noc (E29C N121S) variant, which binds CTP (albeit at a reduced affinity) but cannot hydrolyze CTP, crosslinked similarly to Noc (E29C) (~30% crosslinked fraction) at the saturating concentration of 1 mM CTP, although the *NBS*-stimulated crosslinking was abolished (Figure 2D). Lastly, the NocNΔ10 (E29C) variant, which lacks the N-terminal membrane targeting sequence, crosslinked similarly to Noc (E29C) in the presence of CTP and *NBS* DNA (Figure S2C).

**Figure 2.**
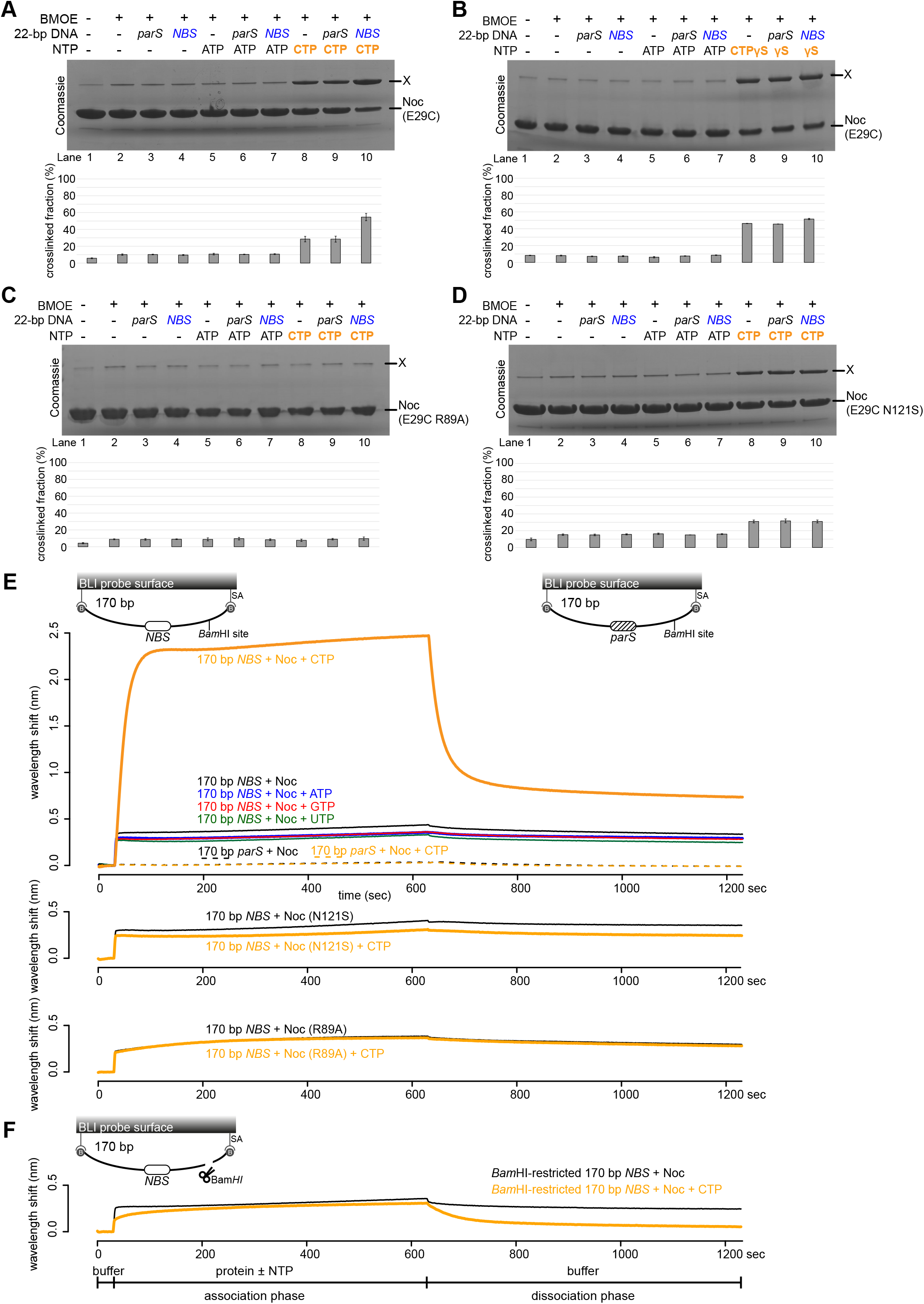
CTP and *NBS* DNA promote the self-engagement of the N-terminal domain of Noc. **(A-B)** SDS-PAGE analysis of BMOE crosslinking products of 10 μM *B. subtilis* Noc (E29C) ± 1 μM 22-bp *parS*/*NBS* DNA ± 1.0 mM NTP. All crosslinking reactions were performed at 22°C unless indicated otherwise. X indicates a crosslinked form of Noc (E29C). Quantification of the crosslinked fraction is shown below each representative image. Error bars represent SEM from three replicates. **(C)** Same as panel A, but Noc (E29C R89A) was used instead. **(D)** Same as panel A, but Noc (E29C N121S) was used instead. **(E)** CTP facilitates the association of Noc with a closed *NBS* DNA substrate beyond nucleation. Bio-layer interferometry (BLI) analysis of the interaction between a premix of 1.0 μM *B. subtilis* Noc ± 1.0 mM NTP and a 170-bp dual biotin-labeled DNA that contains either an *NBS* or a non-cognate *parS* site. Interactions between a dual biotinylated DNA and a streptavidin (SA)-coated probe created a closed DNA molecule where both ends were blocked (Jalal et al., 2020a) (see also the schematic diagram of the BLI probes). Other Noc variants, Noc (R89A) and Noc (N121S), were also analyzed in the same assay. **(F)** BLI analysis of the interaction between a premix of 1.0 μM *B. subtilis* Noc ± 1.0 mM CTP and a BamHI-restricted dual biotinylated *NBS* DNA. The 170-bp *NBS* DNA substrate was also designed with a BamHI recognition site (see the schematic diagram of the probes). The intact dual biotinylated DNA was first immobilized onto the probe surface, then an open end was generated by digestion with BamHI (See STAR Methods). Afterward, the probe was used in a BLI analysis with a premix of Noc ± CTP. Each experiment was triplicated and a representative sensorgram was shown.

### Noc associates with a closed *NBS* DNA substrate in a CTP-dependent manner

To further investigate the roles of the NTD engagement, we followed the spreading of Noc in real-time. We employed a 170-bp dual biotin-labeled *NBS* DNA that had been tethered at both ends onto a streptavidin-coated probe to form a closed DNA, and measured the bio-layer interferometry (BLI) signal (Figure 2E-F) ^9^. BLI monitors wavelength shifts resulting from changes in the optical thickness of the probe during the association/dissociation of Noc from the closed *NBS* DNA substrate. In the absence of CTP, we observed only the nucleation event on *NBS* DNA with 1 μM purified Noc (Figure 2E). Premixing Noc with ATP, GTP, or UTP did not change the sensorgram markedly; however, the addition of 1 mM CTP increased the BLI response by ~sixfold (Figure 2E), consistent with Noc-CTP spreading from the *NBS* to accumulate more on the 170-bp closed DNA substrate than by nucleation alone. We did not observe a noticeable BLI response when a 170-bp closed *parS* DNA substrate was employed instead (Figure 2E), confirming that nucleation and spreading by *B. subtilis* Noc is strictly dependent on the *NBS*. We also observed that DNA-bound Noc-CTP dissociated readily into the solution when the BLI probe was returned to a protein-free buffer without CTP (Figure 2E and Figure S2F, dissociation phase). However, the dissociation of pre-bound Noc-CTP from DNA was slowed by ~fivefold if the probe was returned to a buffer supplemented with CTP (Figure S2F). Furthermore, we observed that pre-bound Noc-CTPγS was more stable and dissociated slowly into a buffer only solution (Figure S2G).

We then tested the mutant proteins by BLI assay and observed that both the Noc (R89A) and Noc (N121S) variants could nucleate but could not spread to accumulate on the closed *NBS* substrate even in the presence of 1 mM CTP (Figure 2E), suggesting that NTD engagement is required for spreading (see also Figure 2C-D).

Next, we investigated whether a DNA substrate with a free end (an open DNA) could also support Noc accumulation in our BLI setup. The 170-bp dual biotin-labeled was designed with a unique *Bam*HI recognition site flanking the *NBS* site (Figure 2F). To generate a free end on the DNA, the DNA-coated probe was immersed in buffer containing *Bam*HI restriction enzyme. Before *Bam*HI digestion, Noc showed an enhanced association on a closed DNA substrate in the presence of CTP (Figure 2E). However, after *Bam*HI digestion, the addition of CTP did not affect the BLI response beyond the nucleation of Noc at the *NBS* (Figure 2F). We reasoned that, similar to the canonical ParB clamp ^9^, Noc spreads but quickly escapes by sliding off a free DNA end. Overall, our BLI analysis support the idea of a clamp-like Noc-CTP that can spread and accumulate on a closed DNA substrate, most likely by entrapping DNA.

### Noc binds liposomes in the presence of CTP

It has been shown previously *in vivo* that Noc possesses membrane-binding activity and that it brings chromosomal DNA to the cell membrane to prevent cell division ^21^. Puzzlingly, however, we could not observe any noticeable association between purified *B. subtilis* Noc and liposomes by a co-sedimentation assay (lanes 1-4 of Figure 3A). We wondered if CTP might be the missing co-factor that activates the membrane-binding activity of Noc. To test this possibility, purified Noc was incubated with liposomes with or without CTP and 22-bp *NBS* DNA, and ultracentrifuged (Figure 3A and Figure S3A). The pellet contained sedimented liposomes and associating protein, while the supernatant contained unbound protein; protein and DNA species from both fractions were analyzed by polyacrylamide gel electrophoresis. We did not observe a significant increase in the amount of Noc in the pellet when CTP alone, CTP and a non-cognate 22-bp *parS* DNA (lanes 5-10 of Figure 3A), or other nucleotides were employed (Figure S3B). However, in the presence of both CTP and a 22-bp *NBS* DNA, ~45% of the Noc protein was detected robustly in the pellet (lane 11-12 of Figure 3A), suggesting that the *in vitro* membrane-binding activity of Noc is CTP- and *NBS*-dependent. *NBS* DNA is most likely required to promote CTP binding and the membrane-binding activity of Noc, rather than to concentrate a large amount of Noc molecules in the vicinity of *NBS*. Supporting this proposition, the short length of a 22-bp *NBS* DNA duplex should allow only a dimer of Noc or Noc-CTP complex to occupy the DNA at a time. Furthermore, nearly all of the 22-bp *NBS* DNA was present in the supernatant (instead of in the pellet) after centrifugation (Figure 3A), most likely because Noc-CTP clamps rapidly escaped the open linear *NBS* DNA (see also Figure 2F). This result suggests that individual Noc-CTP possesses a substantial membrane-binding capability.

**Figure 3.**
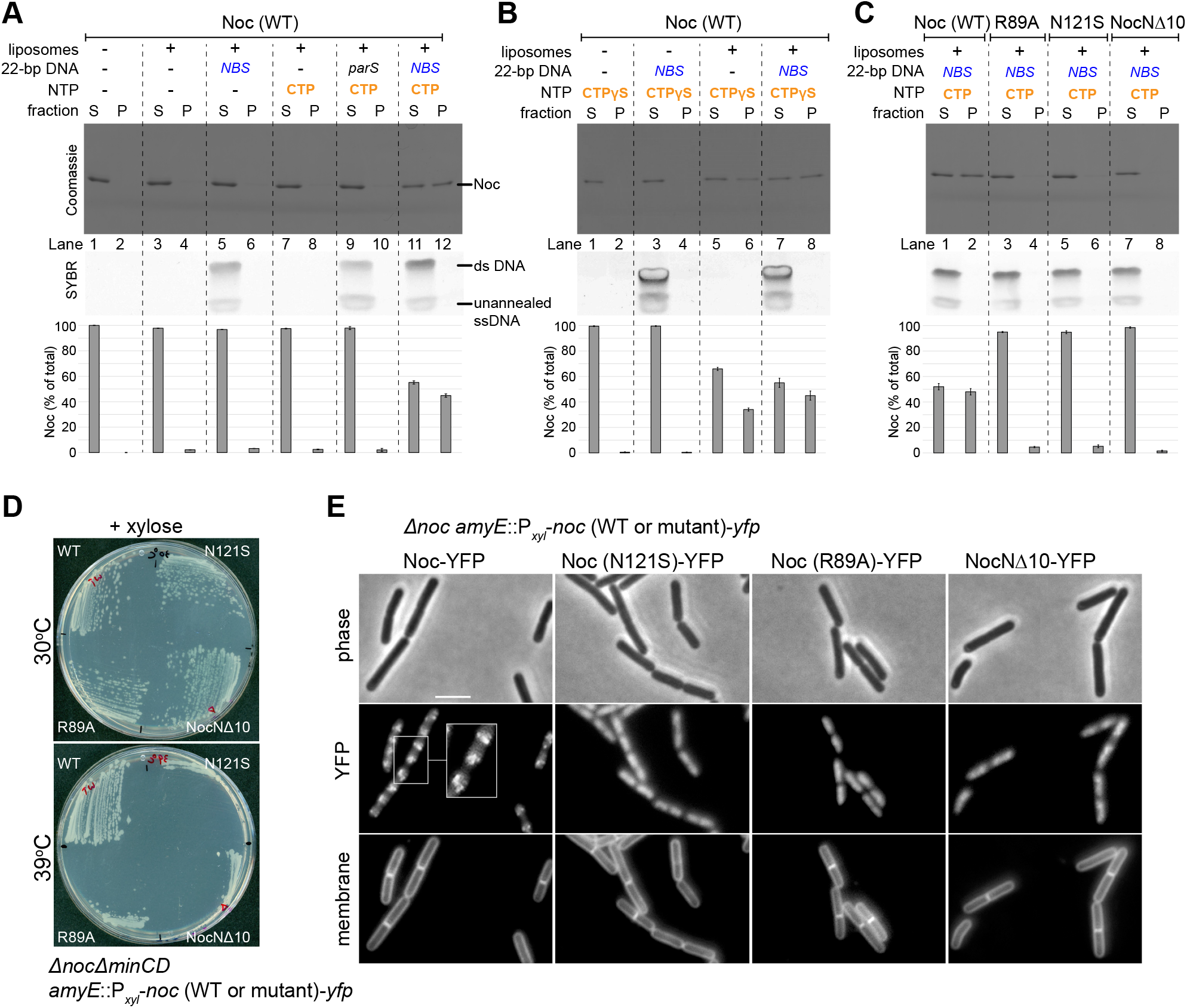
Noc binds liposomes in the presence of CTP and *NBS* DNA, and the phenotypic effects of the Noc variants. **(A)** Analysis of *B. subtilis* Noc binding to membranes by a liposome co-sedimentation assay. A premix of 0.75 μM *B. subtilis* Noc protein ± 1.0 μM 22-bp linear *parS*/*NBS* DNA ± 1.0 mM CTP ± 1.0 mg/mL liposomes was incubated at 22°C for 5 min before ultracentrifugation. The resulting supernatant (S) and pellet (P) fractions were analyzed by SDS-PAGE. Samples were also loaded onto a 20% TBE PAGE, and the gel was subsequently stained with Sybr Green for DNA. Quantification of Noc in each fraction is shown below each representative image. Error bars represent SEM from three replicates. **(B)** Same as panel A but 1.0 mM CTPγS was used instead. **(C)** Other Noc variants, Noc (R89A), Noc (N121S), and NocNΔ10, were also analyzed in a liposome co-sedimentation assay. **(D)** Complementation of *noc* in a *ΔnocΔminCD* background. Strains *ΔnocΔminCD amyE*::P_*xyl*_-*noc* (WT or mutant)-*yfp* were streaked on nutrient agar plates supplemented with 0.5% xylose and incubated at 30°C or 39°C. **(E)** Cellular localization of YFP-labeled Noc (WT or mutants). Representative images of *Δnoc amyE*::P_*xyl*_-*noc* (WT or mutant)-*yfp* cells grown in the presence of 0.5% xylose. Cell membranes were stained with FM5-95. Scale bar, 3 μm. Inset shows a magnification of a section of cells.

Next, we observed that the addition of CTPγS alone caused ~35% of Noc to associate with the pelleted vesicles (lanes 5-6, Figure 3B). The vesicle-bound fraction further increased to ~45% when both CTPγS and a 22-bp *NBS* DNA were present (lanes 7-8, Figure 3B). We infer that CTP binding, but not hydrolysis, is required for the *in vitro* Noc-liposomes interaction. Consistent with the requirement of CTP binding for membrane-binding activity, the Noc (R89A) and Noc (N121S) variants which do not bind CTP or bind CTP weakly failed to co-sediment with liposomes even when CTP and *NBS* were included (lanes 3-4 and 5-6, Figure 3C). The 10-amino-acid N-terminal peptide was previously shown *in vivo* to be the membrane-targeting determinant of *B. subtilis* Noc ^21^. Here, we also confirmed that a purified NocNΔ10 lacking this segment was unable to co-sediment with liposomes *in vitro* regardless of the presence or absence of CTP or *NBS* DNA (lanes 7-8, Figure 3C). Lastly, in another control experiment, *C. crescentus* ParB, which binds CTP but not the cell membrane ^9,10,25^, did not co-sediment with liposomes in the presence of CTP ± *parS* or *NBS* DNA (Figure S3C).

### Noc recruits *NBS* plasmid to liposomes in the presence of CTP

The recruitment of chromosomal DNA to the membrane is essential for Noc to exert the nucleoid occlusion activity *in vivo* ^21^. Indeed, ectopic expression of *noc (R89A)*, *noc (N121S)*, or *nocNΔ10* could not rescue the synthetic cell division defect of a *B. subtilis ΔnocΔminCD* double mutant at elevated temperature (Figure 3D) ^21^. Epi-fluorescence microscopy of *B. subtilis* cells harboring *yfp*-tagged *noc* mutant alleles also confirmed that Noc (R89A), Noc (N121S), and NocNΔ10 failed to form punctate foci near the cell periphery; i.e. were defective in the formation of large membrane-associated nucleoprotein complexes ^21^ (Figure 3E). We wondered if the Noc-dependent recruitment of DNA to the membrane could be biochemically reconstituted. To this end, we assembled a reaction containing purified Noc, liposomes, CTP, and a ~5-kb circular *NBS*-harboring plasmid before ultracentrifugation (Figure 4). Unlike the 22-bp *NBS* DNA, the *NBS* plasmid is topologically closed and therefore should robustly retain closed Noc-CTP clamps. Unfortunately, owing to its high molecular weight, ~45 to 55% of the circular plasmid sedimented independently of the liposomes (lanes 5-6 of Figure 4A, and lanes 1-4 of Figure 4B). Nevertheless, in the presence of liposomes and CTP, the *NBS* plasmid completely co-sedimented with Noc (lanes 11-12 of Figure 4A), demonstrating that Noc can recruit plasmid DNA to liposomes in the presence of CTP. The pellet/supernatant distribution of a control plasmid with no *NBS* (“empty”) was unaffected by the presence of Noc and CTP, and only ~2% of Noc was found in the pellet (lanes 9-10 of Figure 4A). Next, in an attempt to minimize the sedimentation of a plasmid by itself, we performed a vesicle flotation assay in which liposomes and associating protein/DNA migrate up a sucrose gradient to the topmost faction rather than down into the pellet (Figure S4A). Despite the basal level of ~20% total *NBS* plasmid in the top fraction even when liposomes were omitted (lane 3 of Figure S4B), we again observed ~80% of total *NBS* plasmid being recruited to the liposomes when purified Noc and CTP were also present (lane 12 of Figure S4C).

**Figure 4.**
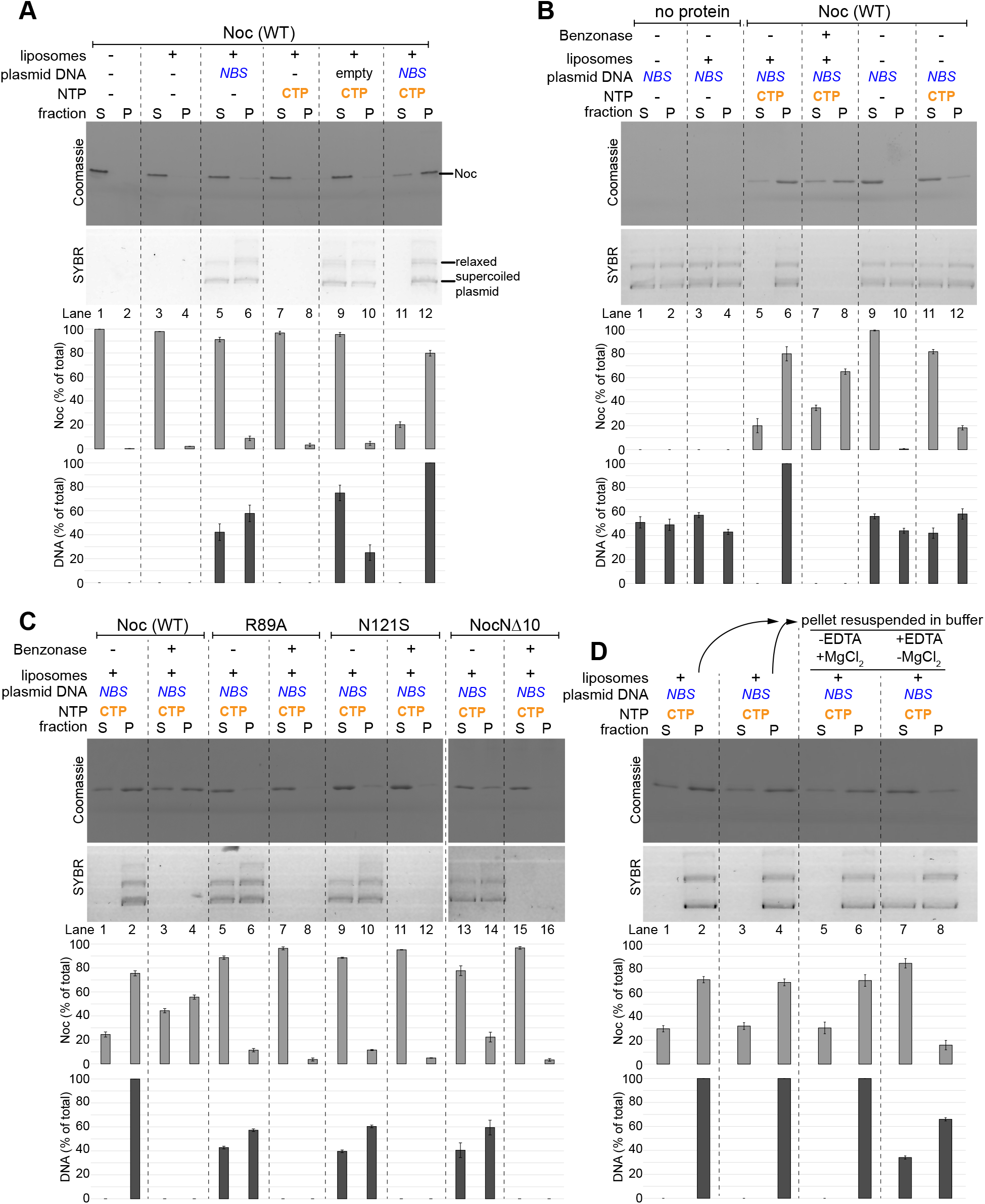
Noc recruits *NBS* plasmid to liposomes in the presence of CTP. **(A)** Analysis of *B. subtilis* Noc binding to membranes and the recruitment of plasmid DNA to the membranes by a liposome co-sedimentation assay. A premix of 0.75 μM *B. subtilis* Noc protein ± 100 nM 5-kb plasmid DNA ± 1.0 mM CTP ± 1.0 mg/mL liposomes was incubated at 22°C for 5 min before ultracentrifugation. Either an empty plasmid or an *NBS*-harboring plasmid was employed in this assay. The resulting supernatant (S) and pellet (P) fractions were analyzed by SDS-PAGE. Samples were also loaded onto a 1% agarose gel, and were subsequently stained with Sybr Green for DNA. Quantification of Noc or DNA in each fraction is shown below each representative image. Error bars represent SEM from three replicates. **(B)** Similar to panel A, a premix of 100 nM *NBS* plasmid ± 0.75 μM Noc ± 1.0 mM CTP ± 1.0 mg/mL liposomes was first assembled and incubated at 22°C for 5 min. However, before ultracentrifugation, a non-specific DNA nuclease (Benzonase) was either added or omitted from the samples, as indicated. **(C)** Other Noc variants, Noc (R89A), Noc (N121S), and NocNΔ10, were also analyzed in a liposome co-sedimentation assay. Benzonase was either added or omitted, as indicated, before ultracentrifugation. **(D)** The association of Noc-*NBS* DNA with liposomes is reversible. A premix of 0.75 μM *B. subtilis* Noc protein + 100 nM *NBS* plasmid + 1.0 mM CTP + 1.0 mg/mL liposomes was ultracentrifuged, and the resulting fractions were analyzed for protein and DNA contents (lanes 1-2 and 3-4). The resulting pellets (lanes 2 and 4) were subsequently resuspended in either a binding buffer (- EDTA + 1 mM MgCl_2_) or in a stripping buffer (+ 10 mM EDTA – MgCl_2_). The resuspensions were ultracentrifuged for the second time, and the resulting fractions were analyzed for protein and DNA contents (lanes 5-6 and 7-8).

In light of the above results, we wondered if the role of *NBS* was to stimulate the membrane-binding activity of Noc-CTP. To test this possibility, we first assembled a co-sedimentation reaction as described above. Subsequently, a non-specific DNase (Benzonase) was added to eliminate the *NBS* plasmid before ultracentrifugation (lanes 7-8 of Figure 4B). The nuclease treatment eliminated intact *NBS* plasmid from both the supernatant and the pellet fractions; however, ~65% of the total amount of Noc still co-sedimented to the pellet in comparison to ~80% when nuclease was omitted (lanes 7-8 vs. lanes 5-6 of Figure 4A). In parallel, we tested spreading-defective Noc (R89A) and Noc (N121S), and a membrane-binding-defective NocNΔ10 for their ability to recruit the *NBS* plasmid to liposomes in a co-sedimentation assay (Figure 4C) as well as in a flotation assay (Figure S4D). Consistent with previous and the above *in vivo* data ^21^, these mutants could not recruit DNA to the pellet fraction beyond the basal level (Figure 4C and Figure S4D). Altogether, these results suggest that the *NBS* specifically activates the membrane-binding activity of Noc in the presence of CTP, thereby recruiting Noc-DNA complexes to the membrane.

### The association of Noc-*NBS* DNA with liposomes is reversible

Once the membrane-associated Noc-DNA nucleoprotein complexes form, can this process be reversed? To investigate, we employed a buffer supplemented with EDTA to sequester Mg^2+^, thereby disrupting CTP binding in preformed liposome-bound Noc-DNA complexes (Figure 4D). In this experiment, the pellet containing preformed liposomes-bound Noc-DNA complexes (lanes 2 or 4 of Figure 4D) was either resuspended in EDTA-minus or EDTA-plus buffer before being ultracentrifuged again. After the second round of centrifugation, both the pellet and the supernatant fractions were analyzed for the presence of protein and DNA (lanes 5-8 of Figure 4D). We observed that while nearly all *NBS* plasmid remained in the pellet when an EDTA-minus buffer was used (lane 6 of Figure 4D), ~38% of the total plasmid returned to the supernatant in the presence of EDTA (lane 7 of Figure 4D). These results demonstrate that the membrane-binding activity of Noc can be reversed, and suggest a possible inhibitory mechanism that keeps apo-Noc in the membrane inactive mode in the absence of CTP.

### Crystal structure of *Geobacillus thermoleovorans* NocΔCTD shows the membrane-targeting amphipathic helix in an autoinhibitory conformation

To gain further insights into the membrane inactive state, we sought to solve a crystal structure of Noc. We could not obtain high-quality crystals of *B. subtilis* Noc either in full-length or truncated forms despite extensive efforts. However, we could grow and collect diffraction data for a C-terminal domain truncated apo-Noc (NocΔCTD) from a thermophilic bacterium, *Geobacillus thermoleovorans*, to 2.5 Å resolution. *B. subtilis* Noc and *G. thermoleovorans* Noc share 72% sequence identity (see the sequence alignment in Figure S5A). The structure was solved by iodide SAD phasing since no other Noc protein family structure was available as a template for molecular replacement. The asymmetric unit contains two similar copies of monomeric apo-Noc (RMSD = 0.9 Å) (Figure S5B), hence we used the more complete subunit for all further analysis.

Each NocΔCTD subunit contains an N-terminal domain (NTD) (helices α1-α6) and an *NBS*-specific DNA-binding domain (DBD) (helices α7-α12) (Figure 5A) ^17,24^. The primary dimerization domain at the C-terminal side of Noc was truncated in the NocΔCTD, and hence was not present in this structure. Most notably, electron density for five of the 10 amino acids (AA) comprising the membrane-targeting helix α1 were visible in a 3_10_ helical conformation (Figure 5A-B). From the structure, it is apparent that the visible membrane-targeting sequence (AA 5 to 10) of Noc is indeed amphipathic, with distinct polar and hydrophobic faces (Figure 5B). The amphipathic helix α1 is immediately followed by helix α2 and subsequently by an 8-AA α2-β1 loop that precedes the main NTD (Figure 5A). By sequence comparison with a canonical ParB ^2^, the main NTD (β1-α6) of Noc contains the CTP-binding motifs, while the amphipathic α1, α2, and the α2-β1 loop are specific to the Noc protein family (Figure S5A). We noted that the hydrophobic face of the amphipathic α1 helix is buried towards α5 and α7 at the core of Noc (Figure 5A and 5C), and thus is unexposed and unlikely to be available for membrane interaction. Specifically, the side chain of S6 hydrogen bonds with the side chain of R150, and the side chain of R7 hydrogen bonds with the main chain oxygen of N104 (Figure 5C). Additionally, the main chain oxygen of S10 hydrogen bonds with the side chain of Q16, and lastly, the main chain oxygen of F11 hydrogen bonds with the side chain of Q120 (Figure 5C). Sidechains of F9 and F11 also interact hydrophobically with the side chains of I116 and I81, respectively (Figure 5C). These interactions thus bury α1 in a potential autoinhibitory conformation. We further noted that α2, which does not target the membrane *per se*, but is conserved in Noc homologs (Figure S5A) ^21^, also contributes to holding α1 in the repressed state (Figure 5D). Specifically, the side chains of both E13 and Q16 form water-mediated contacts with the side chains of I65, R86, and K103, while the side chain of E20 forms a hydrogen bond with Q66 (Figure 5D). Overall, our apo-NocΔCTD structure suggests a repressed state that might keep Noc in the membrane inactive state in the absence of CTP.

**Figure 5.**
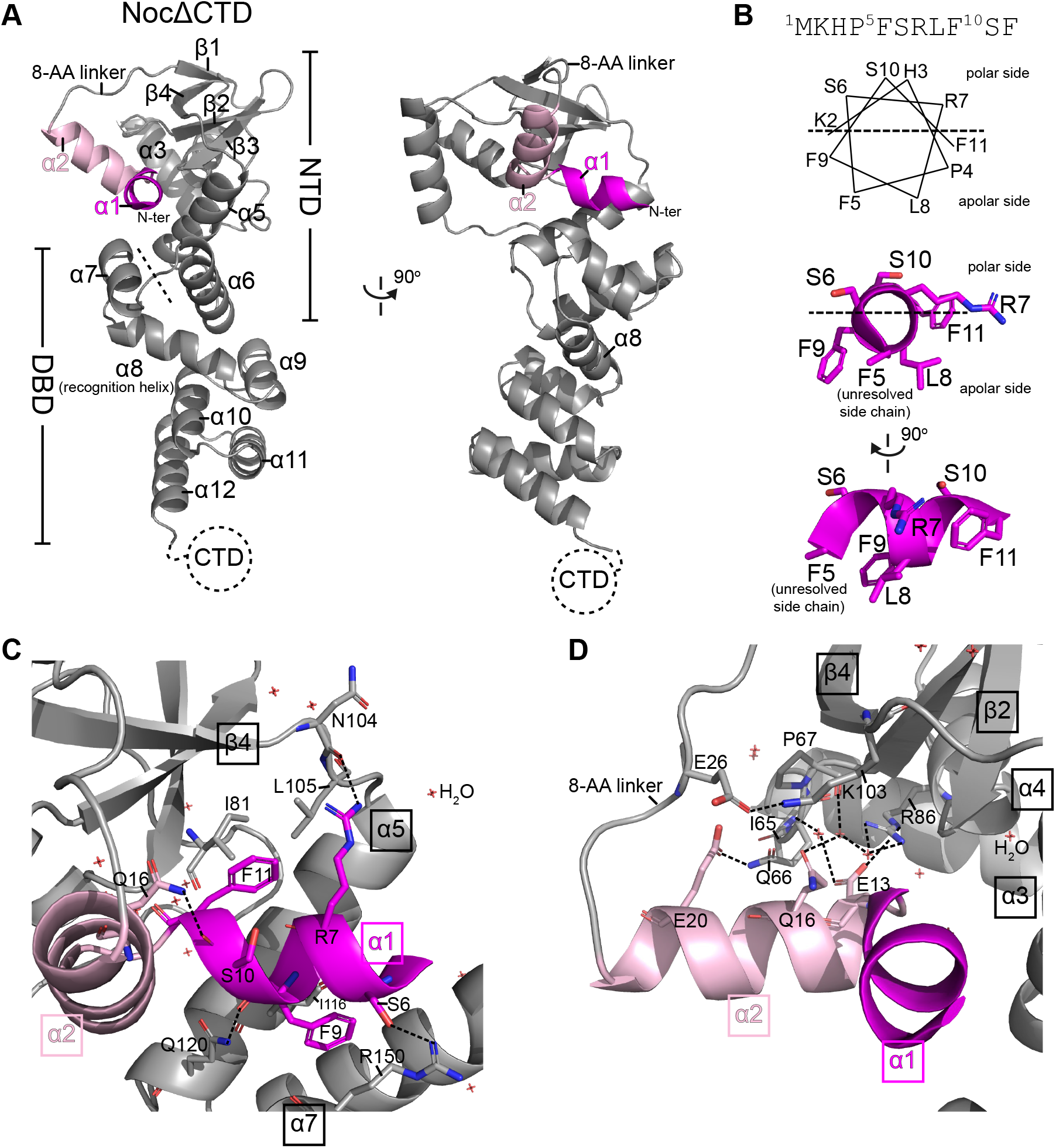
Crystal structure of *Geobacillus thermoleovorans* NocΔCTD shows the membrane-targeting amphipathic helix in an autoinhibitory conformation. **(A)** Crystal structure of a *G. thermoleovorans* NocΔCTD monomer (grey) with an N-terminal amphipathic helix α1 (magenta) and helix α2 (pink). Helix α2 is connected to the main N-terminal domain (NTD) via an eight amino-acid loop. The dashed line demarcates the NTD from the DNA-binding domain (DBD). Helix α8 at the DBD is the recognition helix that contributes to the specific recognition of the *NBS* site (Jalal et al., 2020b; Wu et al., 2009). **(B)** The membrane-targeting amphipathic helix α1. A helical wheel representation of the 10-amino acid at the N-terminus of *G. thermoleovorans* Noc. While the first five amino acids were unresolved in the NocΔCTD crystal structure, the next five amino acids adopt a 310 helical conformation with distinct polar and hydrophobic sides. **(C-D)** Helices α1 (magenta) and α2 (pink) pack themselves into the core N-terminal domain (grey). Hydrogen bonds are shown as dashed lines, water molecules are also shown.

### Crystal structure of the *G. thermoleovorans* NocNΔ26ΔCTD variant is incompatible with an autoinhibitory conformation of the amphipathic helix

Next, we attempted to obtain a co-crystal structure of Noc in complex with nucleotides but were not successful when NocΔCTD or NocNΔ10ΔCTD protein variants were used. However, in the presence of CTPγS, we were able to grow and collect 2.95 Å diffraction data for a crystal of a further truncated *G. thermoleovorans* NocNΔ26ΔCTD variant, which lacks both the N-terminal membrane-targeting helix and the C-terminal domain. After solving its structure, it was apparent that NocNΔ26ΔCTD had adopted an alternative conformation to that of NocΔCTD (Figure 6A vs. Figure 6B-C). This alternative conformation of NocΔN26ΔCTD is compatible with homodimer formation, giving an interfacial area of ~2700 Å^2^ (as evaluated with jsPISA), which resembles that observed for a co-crystal structure of *B. subtilis* ParBΔCTD with bound cytidine diphosphate (CDP) (RMSD = 2.17 Å) (Figure S6A) ^2^. However, there was no clear electron density for a bound nucleotide in our structure; instead a sulfate anion from the crystallization solution occupies a position equivalent to that of the β-phosphate of the nucleotide in the *B. subtilis* ParBΔCTD-CDP structure (Figure S6B) ^2^. We further observed that helix α5 in the NocΔN26ΔCTD structure swings outwards by 104 degrees and no longer forms a bundle with helix α6 from the same subunit (Figure 6A vs. Figure 6B), and that this movement might drive the self-dimerization at the N-terminal domain of Noc, i.e. the NTD engagement (Figure 6C). By superimposing the NTDs of NocΔCTD and NocNΔ26ΔCTD (RMSD = 1.49 Å), we detected severe clashes between α1, α2, and the opposite subunit of NocNΔ26ΔCTD (Figure 6D and Figure S6C-D). Therefore, it is clear that the autoinhibitory state of α1 and α2 (as observed in NocΔCTD) is not compatible with the alternative conformation in the NocNΔ26ΔCTD structure. We speculate that the amphipathic helix α1 and helix α2 might be liberated from the autoinhibitory conformation to be compatible with the NTD-engagement conformation in the NocNΔ26ΔCTD structure.

Lastly, we overexpressed and purified six *B. subtilis* Noc variants to investigate the effects of N-terminal deletions and substitutions on the membrane-binding ability (Figure S7). Removing a lysine residue at position 2 (NocΔK2) or the first four AA (NocΔ4) had a mild effect on the membrane-binding activity of Noc, as judged by liposome co-sedimentation assays (lanes 3-4, and 7-8, Figure S7). However, hydrophobicity-reducing substitutions such as K2E, F5A, or F5E had a negative effect on Noc-liposomes binding (lanes 5-6, lanes 9-10, lanes 11-12, Figure S7). In contrast, when the hydrophobicity of the first 4-AA segment was increased, as in the S4L variant, Noc (S4L) binding to the liposomes increased compared to Noc (WT) (lanes 13-14, Figure S7). Overall, while the conformation of the N-terminal 10-AA sequence of Noc in a membrane-bound state is not yet known, our results suggest that the properties of these amino acids are fine-tuned for membrane-binding activity.

**Figure 6.**
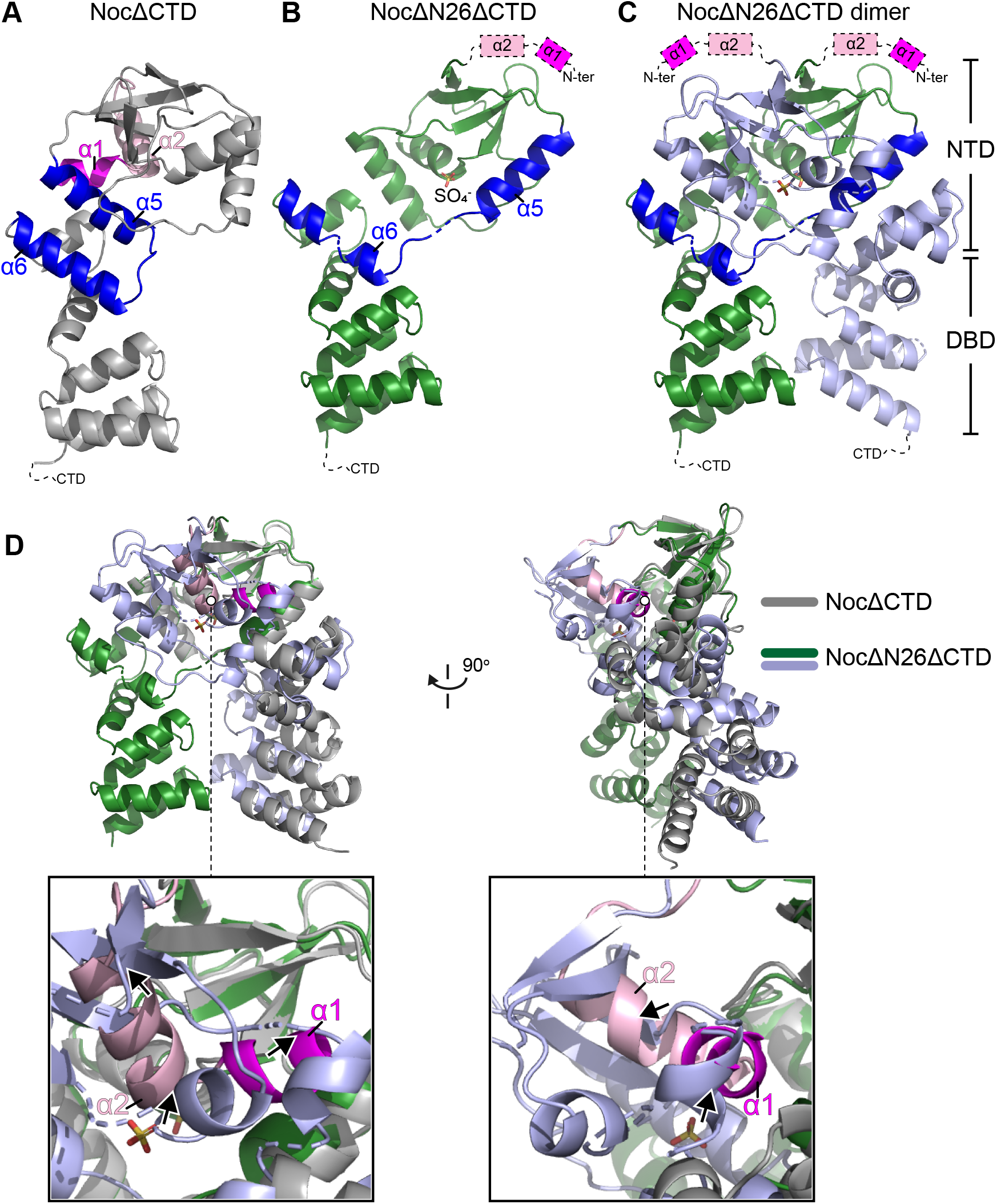
The conformation of Noc in the crystal structure of *G. thermoleovorans* NocNΔ26ΔCTD is incompatible with an autoinhibitory state of the amphipathic helix. **(A)** Crystal structure of *G. thermoleovorans* NocΔCTD (same as Figure 5A) with helices α5 and α6 highlighted in blue. The amphipathic helix α1 and α2 are shown in magenta and pink, respectively. Crystal structure of a *G. thermoleovorans* NocNΔ26ΔCTD variant that lacks both the C-terminal domain (CTD) and helices α1 and α2 (dashed boxes). Helices α5 and α6 are shown in blue. **(C)** A dimer of *G. thermoleovorans* NocNΔ26ΔCTD that self-dimerizes at the N-terminal domain (NTD). Helices α5 and α6 in one of the subunits are shown in blue. **(D)** A superimposition at the NTDs of NocΔCTD monomer (grey) and NocNΔ26ΔCTD dimer (green and light blue) shows a severe clash (arrows) between α1 (magenta), α2 (pink) and the opposite subunit of NocNΔ26ΔCTD (light blue).

## DISCUSSION

### Assembly of the membrane-associated Noc-DNA nucleoprotein complex: roles of CTP

The nucleoid occlusion protein Noc increases cell division efficiency ^22^ by directing the division machinery towards mid-cell either by inhibiting FtsZ formation over the nucleoid ^18,21^ and/or by concentrating FtsZ in the vicinity of a pre-existing mid-cell Z-ring ^23^. The extensive Noc-mediated DNA-membrane interaction is at the heart of both models for nucleoid occlusion ^21,23^. In this study, we show that CTP regulates the nucleoid occlusion activity of Noc. We provide evidence that (i) CTP is required for Noc to form the *NBS*-dependent nucleoprotein complex, and (ii) CTP binding switches Noc from a membrane-inactive auto-repressed state to a membrane-active state. We propose that the dual dependency of Noc’s membrane-binding activity on *NBS* and CTP might ensure productive recruitment of DNA to the bacterial cell membrane to exert the nucleoid occlusion activity (Figure 7). It has been estimated that the intracellular concentrations of Noc and CTP are ~5 μM and ~1 mM, respectively ^24,26^. At these concentrations, if the membrane-binding activity of Noc was solely dependent on CTP, most intracellular DNA-unbound Noc would be in the CTP-bound state and confined to the cell membrane, thus potentially limiting the recruitment of chromosomal DNA to the cell membrane. We therefore reason that the *NBS*-stimulated Noc-CTP interaction provides a mechanism to commit Noc into a pathway in which only DNA-entrapped Noc molecules are able to associate with the cell membrane (Figure 7B). Another consequence of coupling membrane activity to *NBS* binding is that membrane-associated nucleoprotein complexes are spatially confined near the vicinity of *NBS* sites. This spatial confinement is important for directing division machinery formation towards mid-cell where the concentration of *NBS* sites, hence the nucleoid occlusion activity, is lowest ^24^.

**Figure 7.**
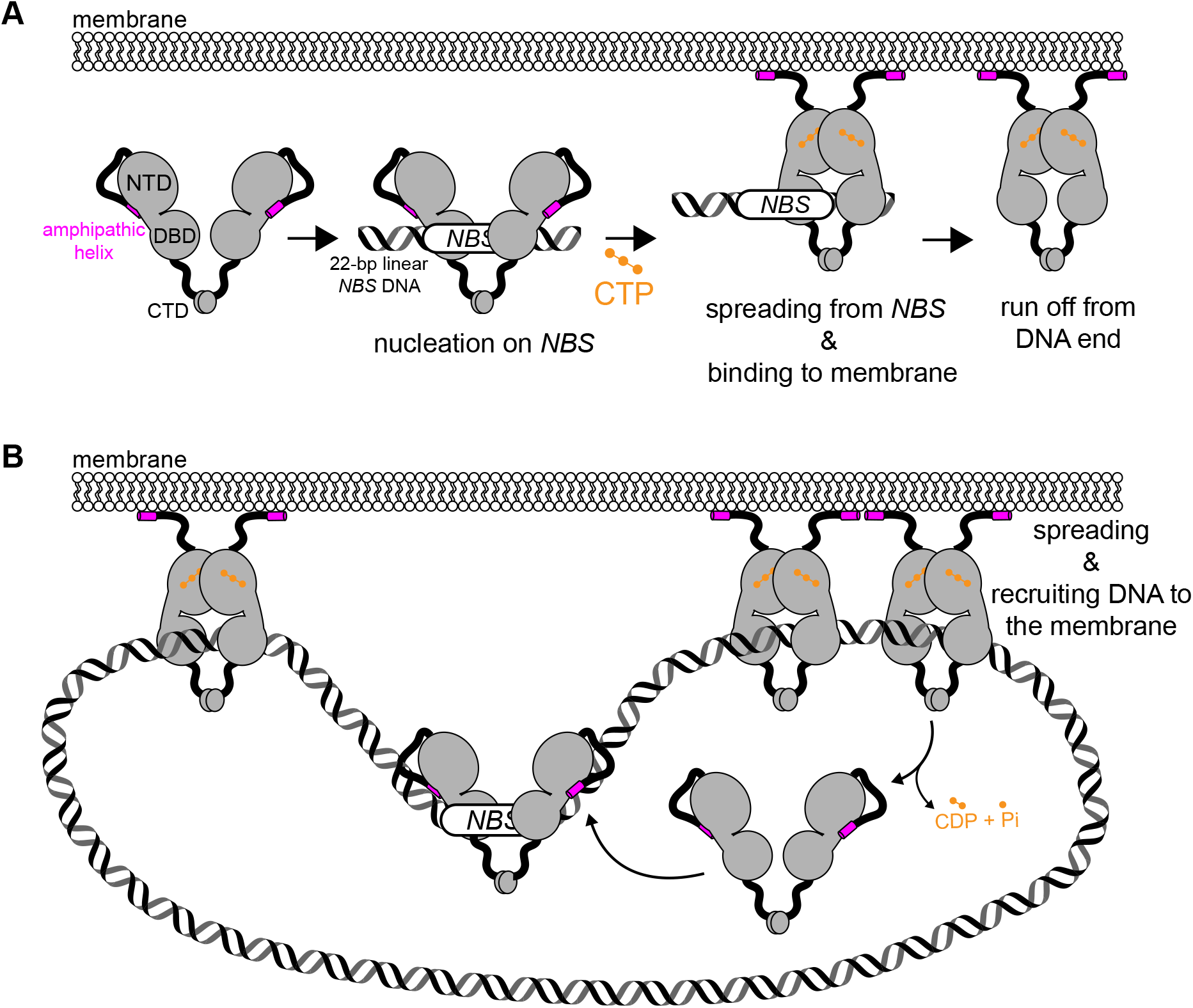
A model for a CTP-dependent regulation of membrane-binding activity of Noc. **(A)** Noc binds specifically to *NBS* site to nucleate on DNA. In the apo- or *NBS*-associated form, the amphipathic helix (magenta) adopts an autoinhibitory conformation, thus cannot bind to the membrane. *NBS*-binding stimulates Noc to bind CTP (orange). Concomitantly, CTP induces a sliding clamp conformation in Noc that can run off open ends of a linear 22-bp *NBS* DNA. In this state, the amphipathic helix is likely liberated from the autoinhibitory conformation, thus enables Noc-CTP to bind to the membrane. **(B)** When a circular *NBS* plasmid with no open end was employed, a sliding clamp of Noc entraps plasmid DNA and recruit DNA to the membrane. In the presence of CTP, a tripartite membrane-protein-DNA linkage is formed. CTP hydrolysis is not required for membrane binding or DNA recruitment but might have a role in releasing Noc from DNA and the membrane.

The lack of a bound nucleotide in the NocNΔ26ΔCTD structure prevents us from drawing a conclusion on the role of CTP and the possible conformational liberation of the amphipathic helix α1. However, several lines of evidence suggest that CTP might favor the NTD-engagement conformation as observed in our NocNΔ26ΔCTD structure: (i) CTP/CTPγS promoted the crosslinking between symmetrical E29C residues, a readout for NTD-engagement (Figure 2A), (ii) the conformation of NocNΔ26ΔCTD is highly similar to that of a nucleotide-bound *B. subtilis* ParB (Figure S6A) ^2^, and (iii) CTP/CTPγS enables Noc (WT) to co-sediment with liposomes (Figure 3). Altogether, it is not unreasonable to speculate that the CTP-induced NTD engagement might liberate the amphipathic helix from its autoinhibitory state to interact with the cell membrane. The 8-AA loop that connects the amphipathic helix to the rest of Noc might offer the flexibility to orient the amphipathic helix parallel to the membrane plane for binding (Figure 7).

This study also provides evidence that *B. subtilis* Noc possesses CTPase activity but CTP hydrolysis is neither required for the clamp formation nor membrane association. What might possibly the role of CTP hydrolysis be? Similar to the counterpart ParB ^1,2,9^, CTP hydrolysis and/or the subsequent release of hydrolytic products might disengage the NTD to open the clamp to release entrapped DNA (Figure 7B). Concomitantly, Noc might revert to the membrane-inactive state, enabling its extraction from the cell membrane. Our experiment that employed EDTA to sequester coordinated Mg^2+^ to artificially promote the dissociation of CTP from Noc supports the proposal that membrane association can be reversible (Figure 4D). Furthermore, *B. subtilis* Noc foci associate with the cell membrane in a transient manner, perhaps suggesting a weak and fast on/off membrane interaction *in vivo* ^21^. The transient association with the membrane, possibly endowed by a CTP hydrolysis event that facilitates the release of Noc, might be advantageous for the cells because a strong and permanent mode of binding might hamper chromosome replication or damage DNA. Supporting this view, fusing NocNΔ10 to a synthetic transmembrane helix led to broken, bisected chromosomes and eventually chromosome segregation defects in many *B. subtilis* cells ^21,27^. Lastly, we noted that NocNΔ10 binds CTP equally well as, or slightly stronger than Noc (WT) (Figure 1D) yet its CTPase rate is reduced by a half (Figure 1E). It is still unclear why this is the case; however, we speculate that the autoinhibitory conformation of this N-terminal-most region might play a role in CTP binding/hydrolysis. Lending support to our speculation, this N-terminal-most region interacts with residues Q66 and R86 (apo-NocΔCTD structure, Figure 5D) whose equivalent residues in *B. subtilis* ParB are known to be critical for CTP binding ^2^.

Overall, we envision a dynamic cycle inside the cells in which CTP hydrolysis converts a closed clamp Noc-CTP to Noc-CDP. The CDP-bound Noc might exist very briefly since CDP likely dissociates rapidly from Noc (due to its low affinity to Noc, Figure S1B), thus opening the clamp to release entrapped Noc from the chromosome and the membrane (Figure 7B). The resulting apo-Noc might also be short-lived because it quickly re-nucleates at *NBS* sites (due to its nM binding affinity to *NBS* ^17^; nucleating Noc at *NBS* then rebinds CTP to close the clamp to spread and to re-bind to the cell membrane, thus essentially restarting the cycle.

### The expanding roles of CTP switches in biology

ATP and GTP switches that control membrane activity are widespread in all domains of life. For example, ATP binding promotes the dimerization of MinD (role in bacterial cell division site selection) and concomitantly increases its affinity for the cell membrane ^28–31^. MinE, a partner of MinD, stimulates the ATPase activity of MinD, promoting MinD dissociation from the membrane ^29,31–33^. In eukaryotes, both the Ras-related protein (Sar) and ADP-ribosylation factor (Arf) (role in vesicle trafficking) function as GTP-dependent switches, cycling between the active GTP-bound form and the inactive GDP-bound form ^34–41^. In the GDP-bound form, the amphipathic helix of Sar/Arf1 adopts a repressed conformation by burying itself into a hydrophobic pocket ^34,42^. The exchange of GDP for GTP induces conformational changes that push the myristoylated amphipathic helix out of the hydrophobic pocket, enabling membrane association ^34,42^. Our study provides the first evidence for a CTP switch that controls membrane-binding activity (Figure 7), suggesting that CTP switches are likely to control more diverse functions in biology than previously appreciated.

## FINAL PERSPECTIVES

Our work unveils a nucleotide-dependent regulatory layer, in addition to the previously described DNA-dependent regulation ^21^, in the activity of the nucleoid occlusion protein Noc. The dual dependency on nucleotide and DNA guarantees a productive formation of the tripartite DNA-protein-membrane super-complex for nucleoid occlusion while allowing efficient cycling of Noc between the membrane-bound and unbound states. In this work, we also provide evidence for a CTP switch that controls membrane-binding activity, adding the control of membrane association in Noc to the role of ParB-CTP in bacterial chromosome segregation. It is likely that CTP switches are pervasive in biology but have so far been underappreciated ^1,2,43^. Finally, evolution is replete with examples of functional domains being adapted to diversify the functions of proteins. The gene encoding Noc apparently resulted from duplication and neo-functionalization of *parB* ^15–17^, our work furthers the understanding of how a CTP switch has been adapted to a new function, and hence might have important implications in understanding biological innovations by evolution.

## Supporting information

PDB files and validation reports

## ACKNOWLEDGEMENTS

This study was funded by the Royal Society University Research Fellowship Renewal (URF\R\201020 to T.B.K.L), and a Wellcome Investigator grant (209500) to Jeff Errington that supported L.J.W. A.S.B.J’s PhD studentship was funded by the Royal Society (RG150448), and N.T.T was funded by the BBSRC grant-in-add (BBS/E/J/000C0683 to the John Innes Centre). We thank Diamond Light Source for access to beamlines I04 and I04-1 under proposals MX18565 and MX25108 with support from the European Community’s Seventh Framework Program (FP7/2007– 2013) under Grant Agreement 283570 (BioStruct-X).

## KEY RESOURCES TABLE

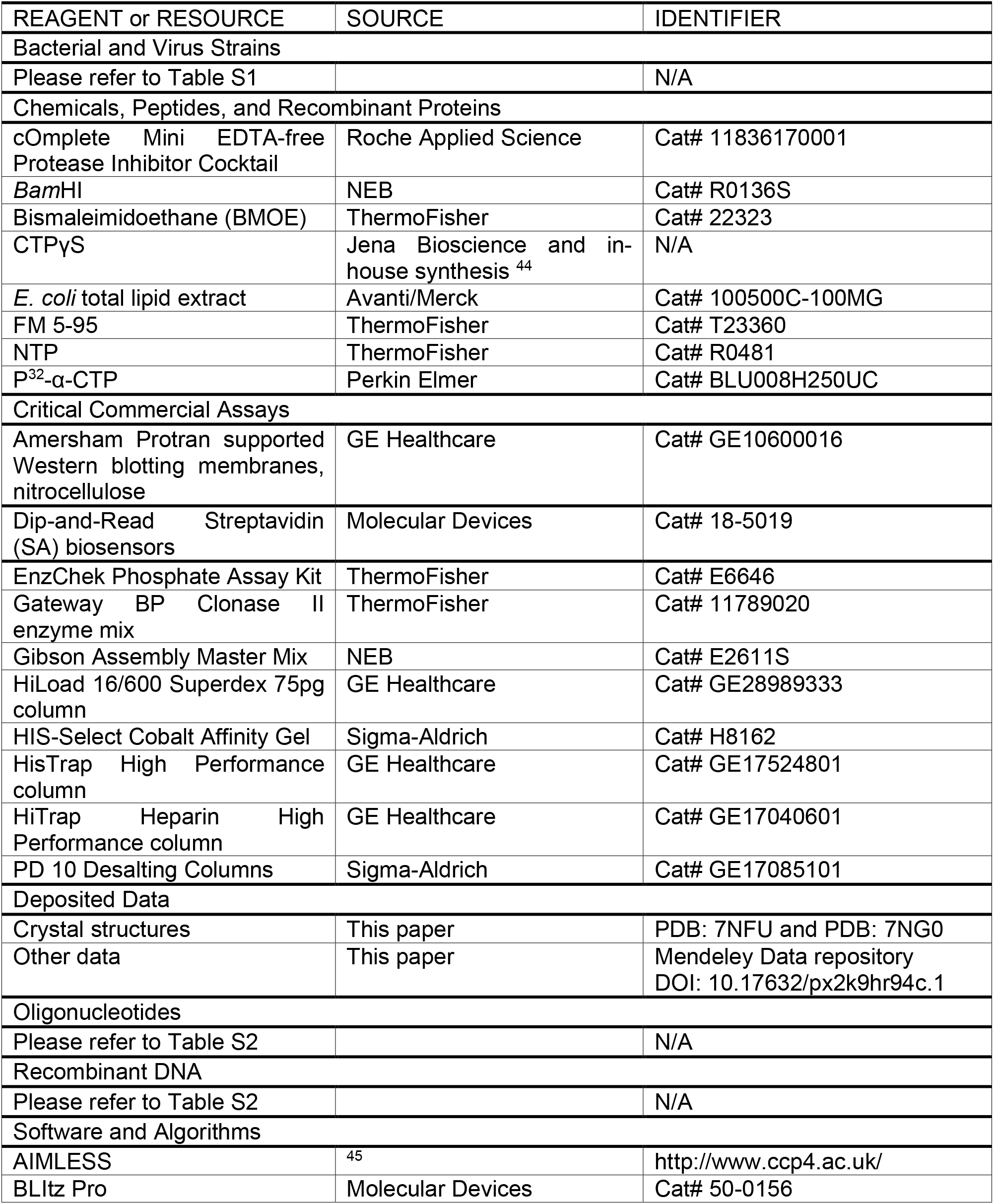

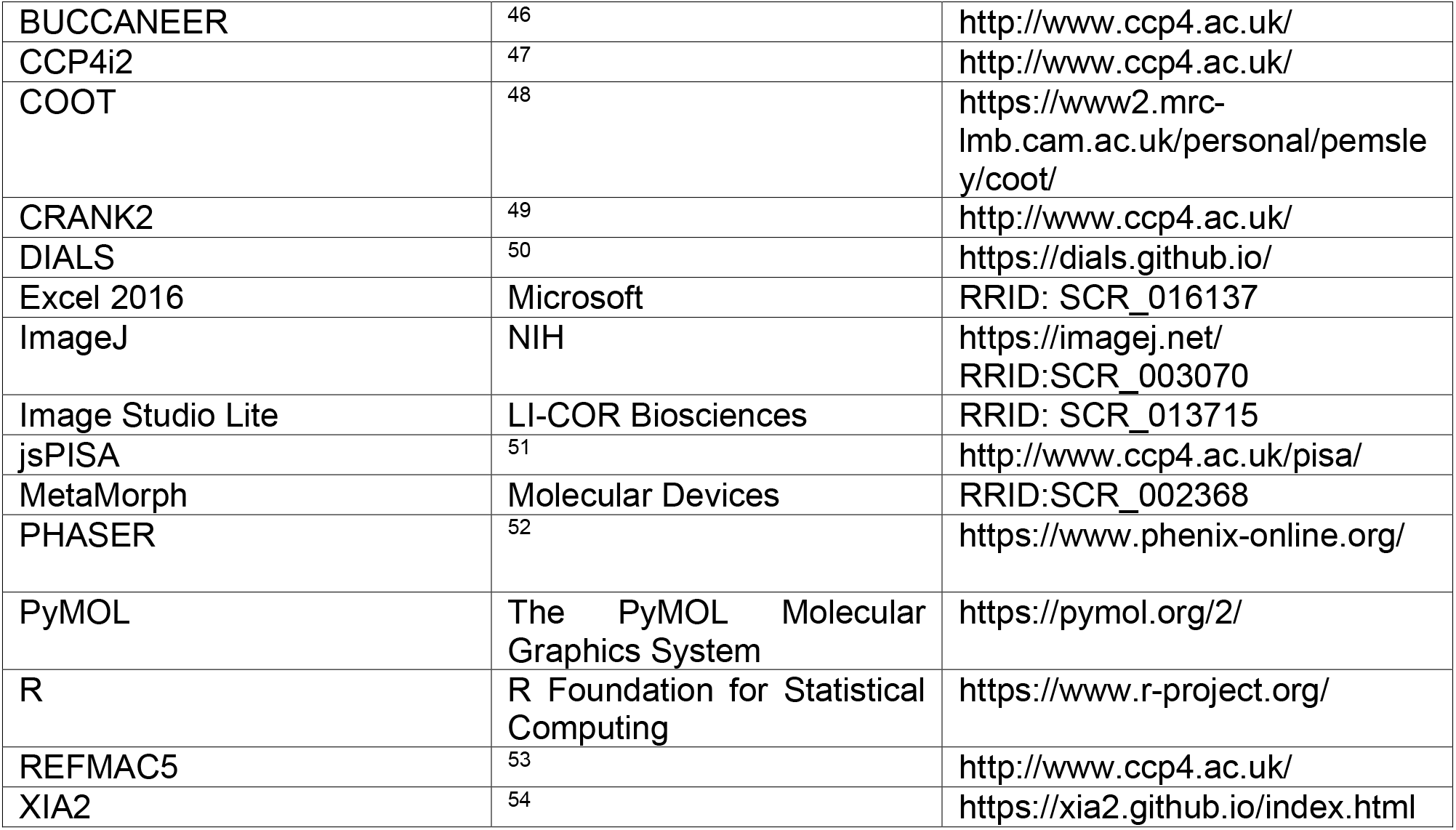

## EXPERIMENTAL MODEL AND METHOD DETAILS

For details about the bacterial strains and culture conditions, please refer to the strain list (Table S1) and Method Details.

### Strains, media, and growth conditions

*Escherichia coli* and *B. subtilis* were grown in LB and CH medium ^55^, respectively. Carbenicillin (100 μg/mL), chloramphenicol (50 μg/mL), and tetracycline (12 μg/mL) were used for selection in *E. coli*, as required. Kanamycin (5 μg/mL), spectinomycin (50 μg/mL), and tetracycline (10 μg/mL) were used for selection in *B. subtilis*, as required. Xylose was added as needed at the concentration indicated.

### Plasmid and strain construction

#### Construction of pET21b::noc (WT)-his_6_

A double-stranded DNA (dsDNA) fragment containing a codon-optimized *B. subtilis noc* gene was chemically synthesized (gBlocks, IDT). The pET21b plasmid backbone was generated via a double digestion of pET21b::*Caulobacter crescentus parB-(his)_6_* with NdeI and HindIII ^10^. The resulting backbone was subsequently gel-purified and assembled with the *noc* gBlocks fragment using a 2x Gibson master mix. Briefly, 2.5 μL of the gBlocks fragment and 2.5 μL of NdeI-HindIII-cut pET21b at equimolar concentration were added to 5 μL of a 2x Gibson master mix (NEB). The mixture was incubated at 50°C for 60 min. Subsequently, 5 μL was used to transform chemically competent *E. coli* DH5α cells. Gibson assembly was possible owing to a 23-bp sequence shared between the NdeI-HindIII-cut pET21b backbone and the gBlocks fragment. The resulting plasmid was verified by Sanger sequencing (Eurofins, Germany).

#### Construction of pET21b::noc (Δ2K, Δ4, K2E, S4L, F5A, or F5E)-his_6_

Same procedure as above, except that dsDNA fragments containing codon-optimized *B. subtilis noc* (Δ2K, Δ4, K2E, S4L, F5A, or F5E) variants were chemically synthesized instead (gBlocks, IDT).

#### Construction of pET21b::nocNΔ10-his_6_

The coding sequence of a 10-amino-acid N-terminally truncated Noc (NocNΔ10) was amplified by PCR using primers AJ65 and AJ66, and pET21b::*noc-his_6_* as a template. The resulting PCR product was gel-purified and assembled into an NdeI-HindIII-cut pET21b using a 2x Gibson master mix. Gibson assembly was possible owing to a 23-bp sequence shared between the NdeI-and-HindIII cut pET21b backbone and the PCR amplified fragment. The 23-bp homologous region was introduced during the synthesis of primers AJ65 and AJ66. The resulting plasmid was verified by Sanger sequencing (Eurofins, Germany).

#### Construction of pET21b::noc (R89A)-his_6_

To introduce the R89A mutation into the coding sequence of Noc, primers P3296 and AJ73, and primers P3297 and AJ74 were used in PCR reactions to amplify the left half and the right half of *noc (R89A)*, respectively, from the *pET21b::noc-his_6_* template. A 15-bp overlapping region between the two PCR fragments contained the point mutation and also enabled their assembly by a Gibson master mix. Briefly, 1.7 μL of each PCR-amplified DNA fragment and 1.6 μL of a gel-purified NdeI-HindIII-cut pET21b at equimolar concentration were added to 5 μL of a 2x Gibson master mix. The mixture was incubated at 50°C for 60 min. Subsequently, 5 μL was used to transform chemically competent *E. coli* DH5α cells. The resulting plasmid was verified by Sanger sequencing (Eurofins, Germany).

#### Construction of pET21b::noc (N121S)-his_6_

The same procedure as above was used to introduce the N121S mutation into the coding sequence of Noc, except that primers P3296 and AJ87, and primers P3297 and AJ86 were used to amplify the left half and the right half of *noc (N121S)*, respectively, from the *pET21b::noc-his_6_* template.

#### Construction of pET21b::noc (E29C)-his_6_

The same procedure as above was used to introduce the E29C mutation into the coding sequence of Noc, except that primers P3296 and AJ85, and primers P3297 and AJ84 were used to amplify the left half and the right half of *noc (E29C)*, respectively, from the *pET21b::noc-his_6_* template.

#### Construction of pET21b::noc (E29C R89A)-his_6_

The same procedure as above was used to introduce the R89A mutation into the coding sequence of Noc (E29C), except that primers P3296 and AJ73, and primers P3297 and AJ74 were used to amplify the left half and the right half of *noc (E29C R89A)*, respectively, from the *pET21b::noc (E29C)-his_6_* template.

#### Construction of pET21b::Noc (E29C N121S)-his_6_

The same procedure as above was used to introduce the N121S mutation into the coding sequence of Noc (E29C), except that primers P3296 and AJ86, and primers P3297 and AJ87 were used to amplify the left half and the right half of *noc (E29C N121S)*, respectively, from the *pET21b::noc (E29C)-his_6_* template.

#### Construction of pET21b::Geobacillus thermoleovorans NocΔCTD-his_6_

A dsDNA fragment containing the coding sequence of a 42-amino-acid C-terminally truncated *G. thermoleovorans* Noc was chemically synthesized (gBlocks, IDT). The gBlocks fragment was assembled into an *Nde*I-*Hin*dIII-cut pET21b using a 2x Gibson master mix. Gibson assembly was possible owing to a 23-bp sequence shared between the NdeI-HindIII-cut pET21b backbone and the gBlocks fragment. The resulting plasmid was verified by Sanger sequencing (Eurofins, Germany).

#### Construction of pET21b::Geobacillus thermoleovorans NocNΔ26ΔCTD-his_6_

The coding sequence of a 26-amino-acid N-terminally truncated and 42-amino-acid C-terminally truncated *G. thermoleovorans* Noc was amplified by PCR using primers AJ76 and AJ81, and pET21b::*Geobacillus thermoleovo*rans NocΔCTD-his_6_ as a template. The resulting PCR product was gel-purified and assembled into an NdeI-HindIII-cut pET21b using a 2x Gibson master mix. Gibson assembly was possible owing to a 23-bp sequence shared between the NdeI-and-HindIII cut pET21b backbone and the PCR amplified fragment. The 23-bp homologous region was introduced during the synthesis of primers AJ76 and AJ81. The resulting plasmid was verified by Sanger sequencing (Eurofins, Germany).

#### Construction of pMCS5-4xNBS

A dsDNA fragment containing four *NBS* sites were chemically synthesized (gBlocks, IDT). The gBlocks fragment was assembled into an EcoRI-cut pMCS5 ^56^ using a 2x Gibson master mix. Gibson assembly was possible owing to a 23-bp sequence shared between the EcoRI-cut pMCS5 backbone and the gBlocks fragment. The resulting plasmid was verified by Sanger sequencing (Eurofins, Germany).

#### Construction of pSG4926N121S and B. subtilis strains harboring the noc(N121S)-yfp fusion

The N121S substitution was introduced into the *noc(WT)-yfp(mut1)* plasmid pSG4926 by site-directed mutagenesis using PFU Turbo DNA polymerase (Agilent Technologies), and primers Noc(N121S)-F and Noc(N121S)-R. After verification of the mutation by DNA sequencing, the mutagenized plasmid (pSG4926N121S) was introduced into the *B. subtilis Δnoc* mutant (DWA117) by transformation ^57^ to generate strain 4746. To construct strain 4747, the *ΔminCminD* deletion was introduced into strain 4746 by transforming strain 4746 with the chromosomal DNA of strain DWA564, with selection for kanamycin resistance. The transformation plates were incubated at 30°C. Note that fusions to YFP(mut1) or mYFP gave a similar localization pattern and were both functional for the wild type Noc.

### DNA preparation for *in vitro* assays

A 22-bp palindromic single-stranded DNA (ssDNA) oligomer (*NBS*: GGA<u>TATTTCCCGGGAAATA</u>TCC or *parS*: GG<u>ATGTTTCACGTGAAACA</u>TCC) [dissolved to 100 μM in buffer containing 1 mM Tris-HCl pH 8.0 and 5 mM NaCl] was heated at 98°C for 5 min before being left to cool down to room temperature overnight to form 50 μM *parS* or *NBS* DNA duplex. The core sequences of *NBS* and *parS* are underlined.

### Construction of biotinylated DNA substrates for bio-layer interferometry (BLI) analysis

DNA constructs were chemically synthesized (gBlocks dsDNA fragments, IDT) with M13F and M13R homologous regions at each end. To generate a dual biotin-labeled DNA substrate, PCR reactions were performed using a 2x GoTaq PCR master mix (Promega), biotin-labeled M13F and biotin-labeled M13R primers, and gBlocks fragments as templates. PCR products were gel purified, and subsequently used in BLI assays.

### Protein overexpression and purification

Plasmids pET21b::*noc-his_6_* (WT or mutants) were introduced into *E. coli* Rosetta (DE3) competent cells by heat-shock transformation. A 10-mL overnight culture was used to inoculate 1 L of LB medium + carbenicillin + chloramphenicol. Cells were grown at 37°C with shaking at 250 rpm to an OD_600_ of ~0.4. The culture was then left to cool down to 4°C before isopropyl-β-D-thiogalactopyranoside (IPTG) was added to a final concentration of 1 mM. The culture was shaken for an additional 3 hrs at 28°C before the cells were pelleted by centrifugation. Pelleted cells were resuspended in a buffer containing 100 mM Tris-HCl pH 8.0, 250 mM NaCl, 10 mM imidazole, 5% (v/v) glycerol, 10 mg of lysozyme, and an EDTA-free protease inhibitor tablet (Merck). Cells were lysed by sonication (10 cycles of 15 s with 10 s resting on ice in between each cycle). The cell debris was removed through centrifugation at 28,000 g for 30 min and the supernatant was filtered through a 0.45 μm filter (Sartorius). The lysate was then loaded into a 1-mL HisTrap column (GE Healthcare) that had been equilibrated with buffer A [100 mM Tris-HCl pH 8.0, 250 mM NaCl, 10 mM imidazole, and 5% (v/v) glycerol]. Protein was eluted from the column using an increasing gradient of imidazole (10 mM to 500 mM) in the same buffer. Noc-containing fractions were pooled together and diluted to a conductivity of 16 mS/cm before being loaded onto a 1-mL Heparin HP column (GE Healthcare) that had been equilibrated with 100 mM Tris-HCl pH 8.0, 25 mM NaCl, and 5% (v/v) glycerol. Protein was eluted from the Heparin column using an increasing gradient of NaCl (25 mM to 1 M) in the same buffer. Lastly, proteins were polished using a gel filtration column. To do so, Noc-containing fractions were concentrated using an Amicon Ultra-15 10 kDa cut-off spin filter (Merck) before being loaded onto a Superdex-75 gel filtration column (GE Healthcare). The gel filtration column was pre-equilibrated with buffer containing 10 mM Tris-HCl pH 8.0 and 250 mM NaCl. Eluted protein fractions were analyzed for purity by SDS-PAGE.

Noc variants that were used in crosslinking experiments were purified using a one-step Ni-affinity column, and all buffers were adjusted to pH 7.4 which was optimal for crosslinking reactions. Purified proteins were desalted using a PD-10 column (Merck), concentrated using an Amicon Ultra-4 10 kDa cut-off spin column (Merck), and stored at −80°C in a storage buffer [100 mM Tris-HCl pH 7.4, 250 mM NaCl, 10% (v/v) glycerol, and 1 mM TCEP].

### Measurement of NTPase activity by EnzChek phosphate assay

NTP hydrolysis was monitored using an EnzChek Phosphate Assay Kit (Thermo Fisher). Samples (100 μL) containing a reaction buffer ± 0 to 1 mM of CTP ± 1 μM *NBS*/*parS* dsDNA + 1 μM of purified Noc (WT or mutants) were assayed in a Biotek EON plate reader for 15 hrs with readings every minute. The reaction buffer (1 mL) typically contained 640 μL ultrapure water, 100 μL 10x customized reaction buffer [100 mM Tris pH 8.0, 2 M NaCl, and 20 mM MgCl_2_], 200 μL MESG substrate solution, and 10 μL purine nucleoside phosphorylase (1 unit). Reactions with buffer only, buffer + protein only or buffer + NTP only were also included as controls. The plates were shaken at 280 rpm continuously for 15 hrs at 25°C. The inorganic phosphate standard curve was also constructed according to the manual. Each assay was triplicated. The NTPase rates were calculated using a linear regression fitting in Excel.

### *In vitro* crosslinking using a sulfhydryl-to-sulfhydryl crosslinker bismaleimidoethane (BMOE)

A 50 μL mixture of 10 μM Noc (WT/mutants) ± 1 mM NTP ± 1 μM 22-bp *NBS*/*parS* dsDNA was assembled in a reaction buffer [10 mM Tris-HCl pH 7.4, 200 mM NaCl, and 1 mM MgCl_2_] and was incubated for 10 min at 22°C or for 1, 5, 10, 15, and 30 min at 4°C. Subsequently, BMOE was added to the final concentration of 1 mM, and the reaction was quickly mixed by three pulses of vortexing. SDS-PAGE sample buffer containing 23 mM β-mercaptoethanol was then added immediately to quench the crosslinking reaction. Samples were heated to 50°C for 10 min before being loaded on 12% WedgeWell Tris-Glycine polyacrylamide gels (Thermo Fisher). Each experiment was triplicated. Polyacrylamide gels were stained in an InstantBlue Coomassie solution (Abcam) and band intensity was quantified using Image Studio Lite (LI-COR Biosciences). The crosslinked fractions were averaged, and their standard errors were calculated in Excel.

CTPγS was custom synthesized either in-house ^44^ or by Jena Biosciences.

### Measurement of protein-DNA interactions by bio-layer interferometry (BLI)

Bio-layer interferometry (BLI) experiments were conducted using a BLItz system equipped with Dip-and-Read Streptavidin (SA) Biosensors (Molecular Devices). BLItz monitors wavelength shifts (nm) resulting from changes in the optical thickness of the sensor surface during association or dissociation of the analyte. All BLI experiments were performed at 22°C. Briefly, the streptavidin biosensor was hydrated in a low-salt binding buffer [100 mM Tris-HCl pH 8.0, 150 mM NaCl, 1 mM MgCl_2_, and 0.005% Tween 20] for at least 10 min before each experiment. Biotinylated double-stranded DNA (dsDNA) was immobilized onto the surface of the SA sensor through a cycle of Baseline (30 s), Association (120 s), and Dissociation (120 s). Briefly, the tip of the biosensor was dipped into a binding buffer for 30 s to establish the baseline, then to 1 μM biotinylated dsDNA for 120 s, and finally to a low salt binding buffer for 120 s to allow for dissociation. After the immobilization of DNA on the sensor, association reactions were monitored at 1 μM dimer concentration of Noc (WT/mutants) (with or without 1mM NTP) for 600 s. At the end of each binding step, the sensor was transferred into a protein-free binding buffer to follow the dissociation kinetics for 600 s. The sensor can be recycled by dipping in a high-salt buffer [100 mM Tris-HCl pH 8.0, 2 M NaCl, 1mM EDTA, and 0.005% Tween 20] for 20 min to remove bound Noc.

For experiments where a closed DNA substrate was cleaved to generate a free DNA end, DNA-coated sensors were dipped into 300 μL of restriction solution [266 μL of water, 30 μL of 10x CutSmart buffer (NEB), and 4 μL of *Bam*HI-HF restriction enzyme (20,000 units/mL)] for 30 min at 37°C.

### Differential radial capillary action of ligand assay (DRaCALA)

Purified Noc (WT and variants) (final concentration: 30 μM) were incubated with 5 nM radiolabeled P^32^-α-CTP (Perkin Elmer), 30 μM of unlabeled CTP (Thermo Fisher), 1 μM of 22 bp *NBS* or *parS* DNA in the reaction buffer [100 mM Tris pH 8.0, 100 mM NaCl, and 5 mM CaCl_2_] for 5 min at room temperature. For the NTP competition assay, the mixture was further supplemented with 500 μM of either unlabeled CTP, CDP, ATP, GTP, or UTP. Four μL of samples were spotted slowly onto a dry nitrocellulose membrane (Amersham Protran 0.45 μm) and air-dried. Subsequently, membranes were exposed to a phosphor screen (GE Healthcare) for two minutes. Each DRaCALA assay was triplicated, and a representative autoradiograph was shown.

### Measurement of Noc-CTPγS/CDP interaction by isothermal titration calorimetry (ITC)

All ITC experiments were recorded using a MicroCal PEAQ ITC instrument (Malvern Panalytical, UK). Experiments were performed at 4°C and both protein and ligand were in the buffer 100 mM Tris-HCl pH 8.0, 100 mM NaCl, and 5 mM CaCl_2_. The calorimetric cell was filled with 100 μM monomer concentration of either *B. subtilis* Noc (WT), Noc (R89A), or Noc (N121S), and was titrated with 3 mM CTPγS. For each ITC run, a single injection of 0.5 μL of 3 mM CTPγS or CDP was performed first, followed by 19 injections of 2 μL each. Injections were carried out at 120 s intervals with a stirring speed of 750 rpm. Each experiment was run in duplicate. The raw titration data were integrated and fitted to a one-site binding model using the built-in software of the MicroCal PEAQ ITC. Controls (CTPγS/CDP into buffer and buffer into protein) were performed and no signal was observed.

### Liposomes preparation

*E. coli* total lipid extract (25 mg/mL in chloroform, Avanti) was used to generate model liposomes. Briefly, an argon stream was used to evaporate chloroform from the lipids, and the resulting lipid cake was further dried under vacuum for 2 hrs. The lipids were subsequently re-suspended in a buffer containing 100 mM Tris-HCl pH 7.4 and 200 mM NaCl. The mixture was incubated at 30°C for 30 min with vigorous vortexing every 5 min. The final concentration of the resuspended lipids was 50 mg/mL. The resuspended lipids were then extruded to ~100 nm single unilamellar vesicles (SUV) using a mini-extruder (Avanti) equipped with polycarbonate membranes (0.1 μm pore size). The size of the resulting SUVs was confirmed by dynamic light scattering.

### Liposome sedimentation assays

A 500 μL mixture of 0.75 μM Noc (WT/mutants) ± 1.0 mg/mL liposomes ± 1 mM NTP ± DNA (either 1 μM of 22-bp *NBS*/*parS* dsDNA or 100 nM of *NBS*-harboring/empty plasmid) was assembled in a binding buffer [100 mM Tris-HCl pH 7.4, 200 mM NaCl, and 1 mM MgCl_2_]. The mixture was incubated at 22°C for 20 min before being centrifuged at 50,000 rpm for 20 min at 22°C (TLA120.2 rotor, Optima Max-E Benchtop Ultracentrifuge). After centrifugation, the supernatant was transferred to a new 1.5-mL Eppendorf tube. The pellet was resuspended in 500 μL of binding buffer before being transferred to another 1.5-mL Eppendorf tube. SDS-PAGE sample buffer was then added, and the samples were heated at 70⁰C for 5 min before being loaded onto either 12% WedgeWell Tris-Glycine polyacrylamide gels, Novex 20% TBE polyacrylamide gels, or 1% agarose gels. Gels were subsequently stained in an InstantBlue Coomassie solution (to detect protein bands) or in a Sybr Green solution (to detect DNA bands). Each assay was triplicated. Protein/DNA band intensity was quantified using Image Studio Lite (LI-COR Biosciences). The protein/DNA fractions were averaged, and their standard errors were calculated in Excel.

For nuclease treatment, a 500 μL mixture of 0.75 μM Noc (WT/mutants) ± 1.0 mg/mL liposomes ± 1 mM NTP ± 100 nM of *NBS*-harboring/empty plasmid was incubated at room temperature for 10 min. Afterward, 1 μL of Benzonase (250 units/ μL) was added, and the mixture was incubated for a further 10 min at 22°C before ultracentrifugation.

For re-sedimentation experiments (Figure 4D), the pellet (from the first round of ultracentrifugation) was resuspended either in 500 μL of binding buffer [100 mM Tris-HCl pH 7.4, 200 mM NaCl, and 1 mM MgCl_2_] or in a stripping buffer [100 mM Tris-HCl pH 7.4, 200 mM NaCl, and 10 mM EDTA]. The resuspended pellet was centrifuged for the second time at 50,000 rpm for 20 min at 22°C. After the second centrifugation, the supernatant was transferred to a new 1.5-mL Eppendorf tube. The pellet was resuspended in 500 μL of binding buffer before being transferred to another 1.5-mL Eppendorf tube.

SDS-PAGE sample buffer was then added to the supernatant and the pellet fractions, and the samples were analyzed on denaturing polyacrylamide gels.

### Liposome flotation assays

A 200 μL mixture of 0.75 μM Noc (WT/mutants) ± 1.0 mg/mL liposomes ± 1 mM NTP ± 20 nM *NBS*-harboring/empty plasmid was assembled in a 30% sucrose buffer [100 mM Tris-HCl pH 7.4, 200 mM NaCl, 1 mM MgCl_2_, and 30% sucrose]. The mixture was incubated at 22°C for 5 min before being overlaid with 250 μL of a 25% sucrose buffer [100 mM Tris-HCl pH 7.4, 200 mM NaCl, 1 mM MgCl_2_, and 25% sucrose]. Finally, 150 μL of a 0% sucrose buffer [100 mM Tris-HCl pH 7.4, 200 mM NaCl, and 1 mM MgCl_2_] was added as the top layer. The solution was incubated for 15 min at 22°C before being centrifuged at 70,000 rpm at 22°C for 20 min (TLA120.2 rotor, Optima Max-E Benchtop Ultracentrifuge). After centrifugation, three equal fractions (200 μL each) were gently drawn sequentially from the bottom of the ultracentrifugation tube using a Hamilton syringe. SDS-PAGE sample buffer was added to each fraction, and samples were heated at 70⁰C for 5 min before being loaded onto either 12% WedgeWell Tris-Glycine polyacrylamide gels or 1% agarose gels. Gels were subsequently stained in an InstantBlue Coomassie solution (to detect protein bands) or in a Sybr Green solution (to detect DNA bands). Each assay was triplicated. Protein/DNA band intensity was quantified using Image Studio Lite (LI-COR Biosciences). The protein/DNA fractions were averaged, and their standard errors were calculated in Excel.

### Protein crystallization, structure determination, and refinement

Crystallization screens were set up in sitting-drop vapor diffusion format in MRC2 96-well crystallization plates with drops comprised of 0.3 μL precipitant solution and 0.3 μL of protein and incubated at 293 K. After optimization of initial hits, suitable crystals were cryoprotected and mounted in Litholoops (Molecular Dimensions) before flash-cooling by plunging into liquid nitrogen. X-ray data were recorded either on beamline I04 or I04-1 at the Diamond Light Source (Oxfordshire, UK) using either an Eiger2 XE 16M or a Pilatus 6M-F hybrid photon counting detector (Dectris), with crystals maintained at 100 K by a Cryojet cryocooler (Oxford Instruments). Diffraction data were integrated and scaled using DIALS ^50^ via the XIA2 expert system ^54^ then merged using AIMLESS ^45^. Data collection statistics are summarized in Table S3. The majority of the downstream analysis was performed through the CCP4i2 graphical user interface ^47^.

#### Geobacillus thermoleovorans NocΔCTD Noc

His-tagged NocΔCTD Noc (~10 mg/mL) was premixed with 1 mM CTP and 1 mM MgCl_2_ in buffer [10 mM Tris-HCl pH 8.0 and 250 mM NaCl] before crystallization. Crystals grew in a solution containing 2.0 M ammonium sulfate and 50 mM tri-sodium citrate, and were cryoprotected in the crystallization solution supplemented with 20% (v/v) glycerol, 1 mM CTP and 1 mM MgCl_2_. For iodide derivatization, the cryoprotectant comprised the crystallization solution supplemented with 25% (v/v) ethylene glycol, 1 mM CTP, 1 mM MgCl_2_, and 500 mM potassium iodide; crystals were soaked in this solution for less than 30 seconds before cryo-cooling. Three 360° passes of X-ray data were taken at a wavelength of 1.8 Å from different parts of a single crystal and merged to give a highly redundant dataset to 3.4 Å resolution in space group *P*2_1_3 with cell parameters *a* = *b* = *c* = 136.6 Å. Solvent content estimation gave a value of 66% for two copies of the 29 kDa subunit per asymmetric unit (ASU). The structure was subsequently solved by single-wavelength anomalous dispersion using the CRANK2 pipeline ^49^, which located 12 iodide sites, and BUCCANEER ^46^ was able to build and sequence 339 residues in two chains corresponding to 67% of those expected for two monomers, giving *R*_work_ and *R*_free_ values of 0.318 and 0.362, respectively, to 3.4 Å resolution after refinement with REFMAC5 ^53^. At this point, this preliminary model was used as a starting point for refinement against a native data set processed to 2.5 Å resolution in the same space group, but with a significantly longer cell edge of 146.8 Å corresponding to a solvent content of 72.6%. Thus, it was necessary to resolve the structure by molecular replacement using PHASER ^52^ before further refinement in REFMAC5. After a complete rebuild in BUCCANEER, several iterations of model building in COOT ^48^ and REFMAC5 refinement jobs yielded the final model with *R*_work_ and *R*_free_ values of 0.210 and 0.240, respectively, to 2.5 Å resolution.

Despite the presence of CTP in the crystallization buffer, no density for CTP was found in the NocΔCTD structure, presumably because *NBS* DNA was not included to facilitate CTP-binding, or the high concentrations of sulfate (2M) from the precipitant excluded the ligand.

### Geobacillus thermoleovorans NocNΔ26ΔCTD

His-tagged NocNΔ26ΔCTD Noc (~10 mg/mL) was premixed with 1 mM CTPγS and 1 mM MgCl_2_ in buffer [10 mM Tris-HCl pH 8.0 and 250 mM NaCl] before crystallization. Crystals grew in a solution containing 0.2 M di-ammonium phosphate and 2.3 M ammonium sulfate and were cryoprotected in this solution supplemented with 20% (v/v) glycerol. X-ray data were processed to a resolution of 2.95 Å in space group *C*222_1_ with cell parameters of *a* = 105.1, *b* = 106.6, *c* = 42.2 Å. Analysis of the likely composition of the ASU suggested that it contained a single copy of the 26 kDa NocNΔ26ΔCTD Noc monomer, giving an estimated solvent content of 46%. The structure was solved by molecular replacement with PHASER ^52^ using chain A from the *G. thermoleovorans* NocΔCTD structure above as the template. The search model was split into three separate ensembles comprising residues 26-100, 101-140, and 141-230, respectively. PHASER successfully placed all three ensembles, although one of these had to be interchanged with a symmetry mate to restore the connectivity of the starting template. Several iterations of model building in COOT and refinement REFMAC5 yielded the final model with *R*_work_ and *R*_free_ values of 0.267 and 0.288, respectively, to 2.95 Å resolution.

### Fluorescence microscopy

Cells containing fluorescent protein fusions were grown at 30°C. Xylose (0.5% w/v) was included in the media to induce the expression of YFP fusions in *B. subtilis*. Cell membranes were stained by mixing 15 μL of culture with 0.5 μL of membrane dye FM5‐95 (200 μg/ml, Invitrogen). Cells were mounted on microscope slides covered with a thin agarose pad (1.2% w/v in dH_2_O) and images were acquired with a Rolera EM-C2 (Q-imaging) camera attached to a Nikon Ti microscope using METAMORPH version 6 (Molecular Devices), with an exposure time of 400 ms for YFP and 1000 ms for membranes. Images were prepared for publication using ImageJ.

## QUANTIFICATION AND STATISTICAL ANALYSIS

Information about statistical analysis and sample size for each experiment are detailed in the relevant methods sections.

---

**Figure S1.**
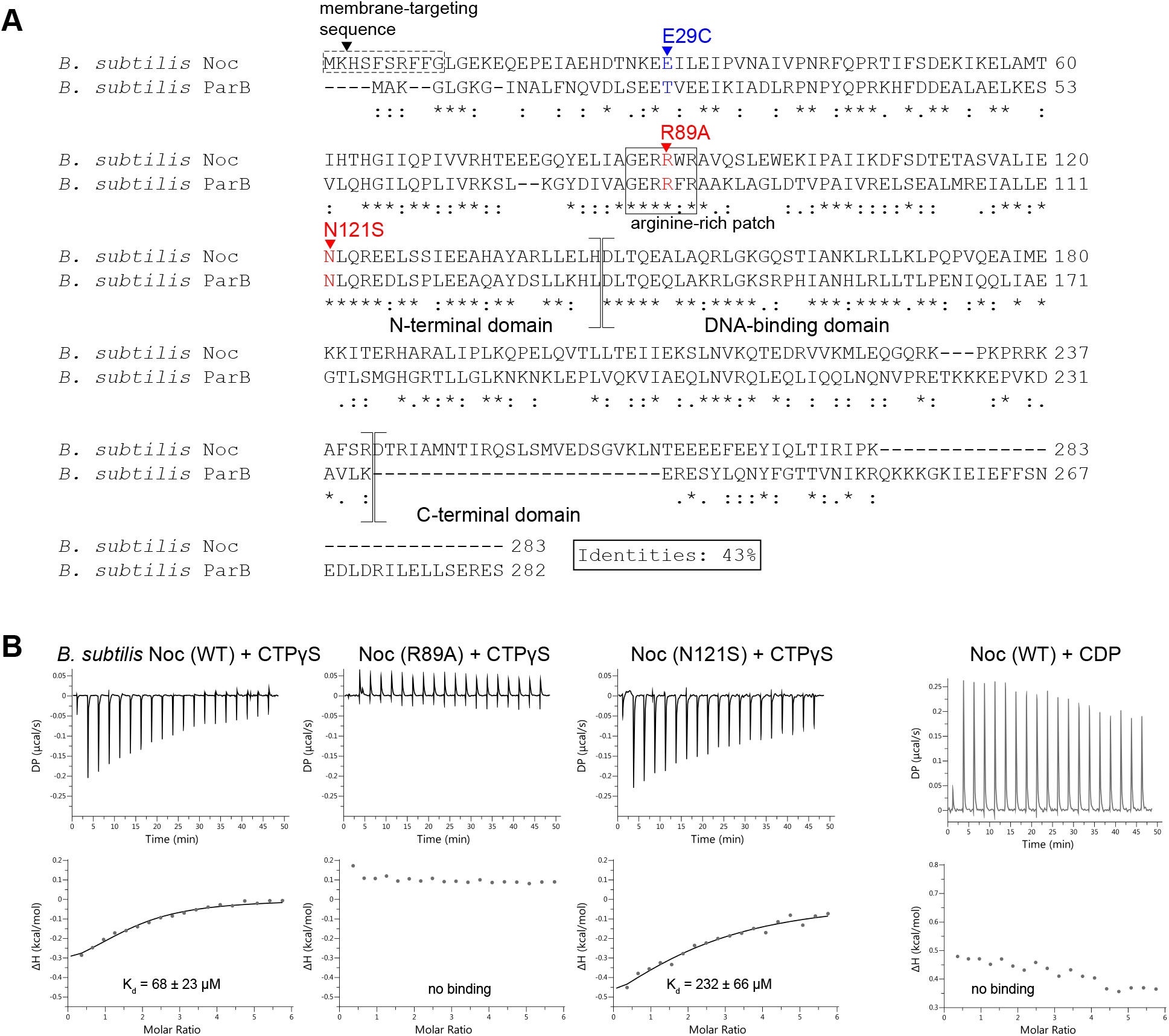
Interactions between Noc (WT and variants) and CTPγS or CDP, related to Figure 1. **(A)** A sequence alignment between *B. subtilis* Noc and its paralog ParB. Residues R89 (red) and N121 (red) in *B. subtilis* Noc, whose equivalent substitutions in *B. subtilis* ParB have been shown to impair spreading and CTP binding (Soh et al., 2019), were substituted by alanine and serine, respectively. Residue E29 (blue) was substituted by cysteine to generate a Noc (E29C) variant which was subsequently used in BMOE crosslinking assays (Figure 2A-D). An equivalent substitution T22C was previously used in BMOE crosslinking assays for in *B. subtilis* ParB (Soh et al., 2019). The conserved arginine-rich patch that mediates CTP binding in *B. subtilis* ParB is shown in a solid box. The 10-amino-acid membrane-targeting sequence of Noc is shown in a dashed box. *B. subtilis* ParB does not possess an equivalent membrane-targeting sequence. The positions of the N-terminal domain, the DNA-binding domain, and the C-terminal domain of Noc and ParB are also indicated on the sequence alignment. **(B)** Analysis of the interaction of *B. subtilis* Noc (WT and mutants) with CTPγS or CDP by isothermal calorimetry (ITC). ITC directly measures the heat released or absorbed during a biomolecular binding event. The large heat exchange from Noc-*NBS* DNA binding (nM binding affinity (Jalal et al., 2020b)) might mask the weaker heat signal from Noc-CTPγS/CDP binding, therefore *NBS* DNA was not included in these ITC experiments. Each experiment was duplicated. Regression curves were fitted, and binding affinities (K_d_) are shown.

**Figure S2.**
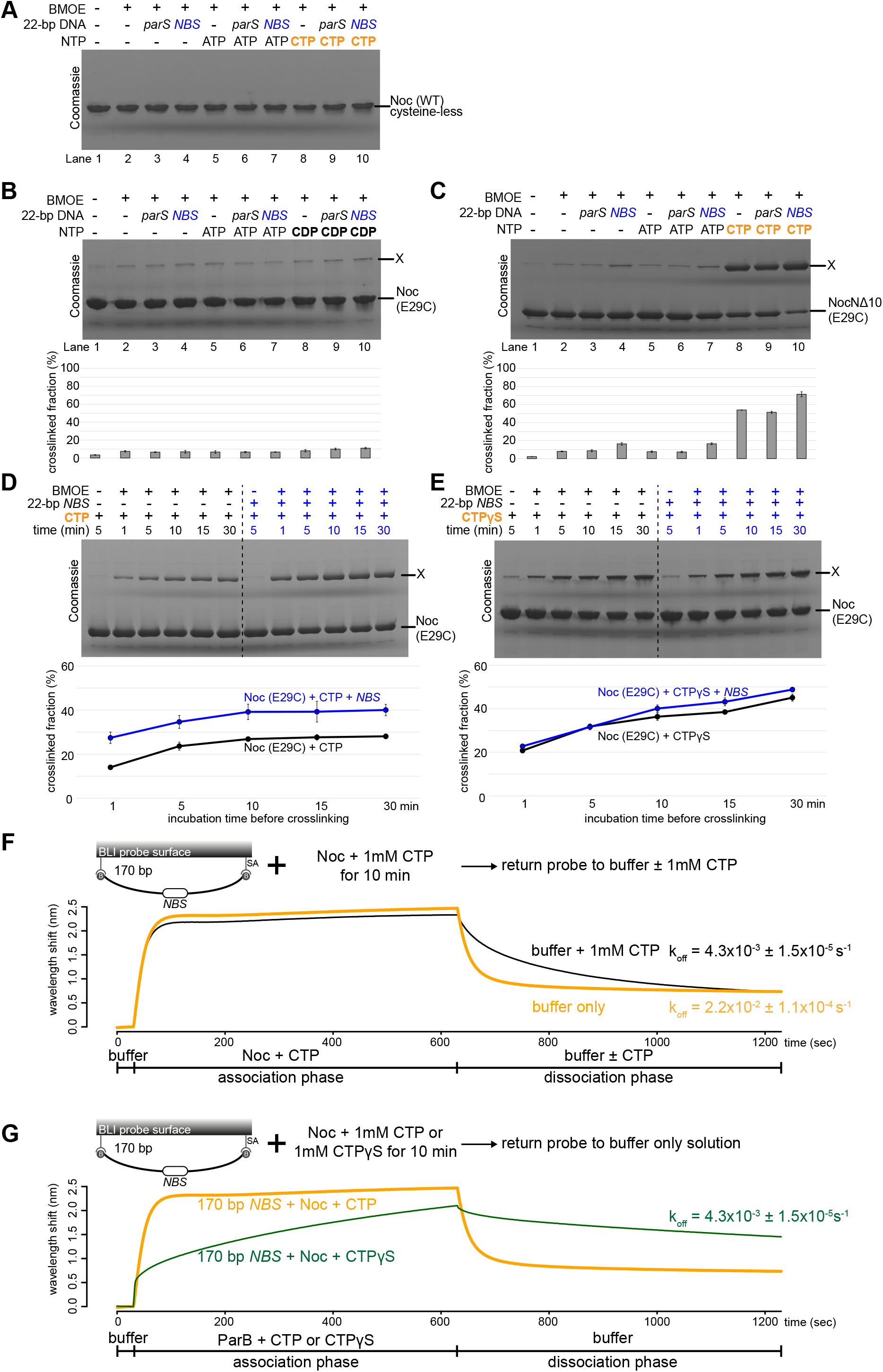
CTP and CTPγS, but not other nucleotides, promote the engagement of the N-terminal domain of Noc, related to Figure 2. **(A)** SDS-PAGE analysis of BMOE crosslinking products of 10 μM *B. subtilis* Noc (WT) protein ± 1.0 μM 22-bp *parS*/*NBS* DNA ± 1.0 mM NTP. Wild-type Noc naturally lacks cysteine, hence does not crosslink in the presence of BMOE. All crosslinking reactions were performed at 22°C unless indicated otherwise. **(B)** Same as panel A but Noc (E29C) was used instead. X indicates a crosslinked form of Noc (E29C). Quantification of the crosslinked fraction is shown below each representative image. Error bars represent SEM from three replicates. Same as panel A, but NocNΔ10 (E29C) was used instead. **(D)** Time-course of Noc (E29C) crosslinking with CTP in the presence or absence of 22-bp *NBS* DNA. Purified Noc (E29C) was preincubated with 1.0 mM CTP ± 1 μM 22-bp *NBS* DNA for 1, 5, 10, 15, or 30 min at 4°C (instead of the usual 22°C) before BMOE was added. A lower incubation temperature was needed to slow down the reaction. Quantification of the crosslinked fraction is shown below each representative image. Error bars represent SE from three replicates. **(E)** Same as panel D, but 1.0 mM CTPγS was used instead. **(F)** BLI analysis of the interaction between *B. subtilis* Noc-CTP and a 170-bp dual biotin-labeled *NBS* DNA. For the association phase, the interaction between a premix of 1.0 μM *B. subtilis* Noc ± 1.0 mM CTP and a 170-bp dual biotin-labeled *NBS* probe was monitored in real-time for 10 min. For the dissociation phase, the probe was returned to either a buffer-only solution or a buffer supplemented with 1mM CTP. The dissociation rate (k_off_) of bound Noc into buffer is shown for each reaction. **(G)** BLI analysis of the interaction between *B. subtilis* Noc-CTP or Noc-CTPγS and a 170-bp dual biotin-labeled *NBS* DNA. For the association phase, the interaction between a premix of 1.0 μM *B. subtilis* Noc + 1.0 mM CTP or CTPγS and a 170-bp dual biotin-labeled *NBS* probe was monitored in real-time for 10 min. For the dissociation phase, the probe was returned to a buffer-only solution. The dissociation rate (k_off_) of bound Noc into buffer is shown for each reaction. Each experiment was triplicated and a representative sensorgram was shown.

**Figure S3.**
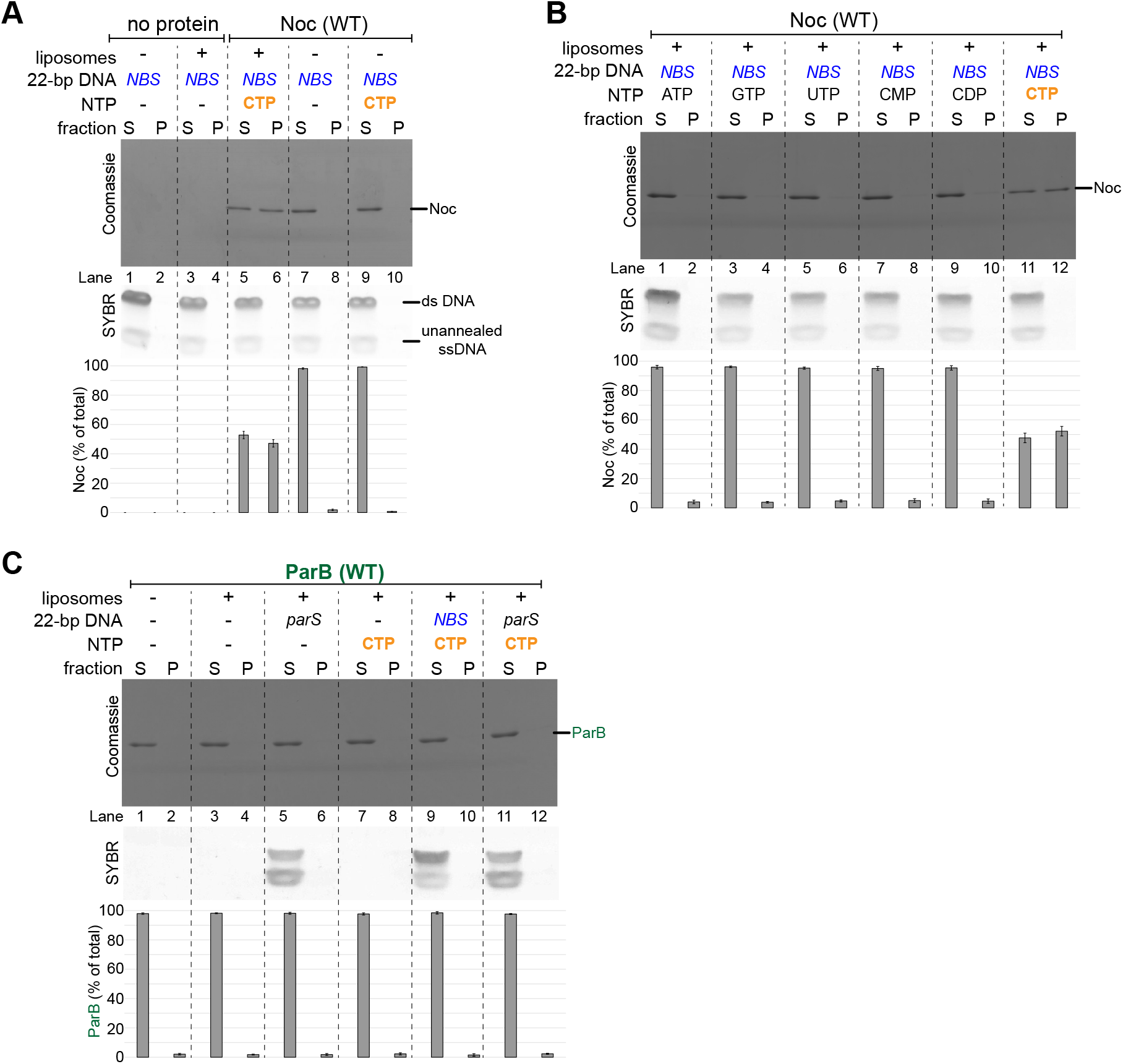
CTP and *NBS* DNA enable Noc binding to liposomes, related to Figure 3. **(A)** Analysis of *B. subtilis* Noc binding to membranes by a liposome co-sedimentation assay. A premix of 1.0 μM 22-bp linear *NBS* DNA ± 0.75 μM *B. subtilis* Noc protein ± 1.0 mM CTP ± 1.0 mg/mL liposomes was incubated at 22°C before ultracentrifugation. The resulting supernatant (S) and pellet (P) fractions were analyzed by SDS-PAGE. Without Noc, 22-bp *NBS* DNA did not co-sediment on its own (lanes 1-2) or with liposomes (lanes 3-4). Samples were also loaded onto a 20% TBE PAGE, and the gel was subsequently stained with Sybr Green for DNA. Quantification of Noc in each fraction is shown below each representative image. Error bars represent SEM from three replicates. **(B)** CTP but no other nucleotide enables Noc to co-sediment with liposomes. A premix of 0.75 μM *B. subtilis* Noc protein + 1.0 μM 22-bp *NBS* DNA ± 1.0 mM NTP + 1.0 mg/mL liposomes was incubated at 22°C for 5 min before ultracentrifugation. The resulting supernatant (S) and pellet (P) fractions were analyzed by SDS-PAGE. **(C)***Caulobacter crescentus* ParB does not co-sediment with liposomes in any tested conditions. A premix of 0.75 μM *C. crescentus* ParB protein ± 1.0 μM 22-bp linear *parS*/*NBS* DNA ± 1.0 mM CTP ± 1.0 mg/mL liposomes was incubated at 22°C for 5 min before ultracentrifugation. The resulting supernatant (S) and pellet (P) fractions were analyzed by SDS-PAGE.

**Figure S4.**
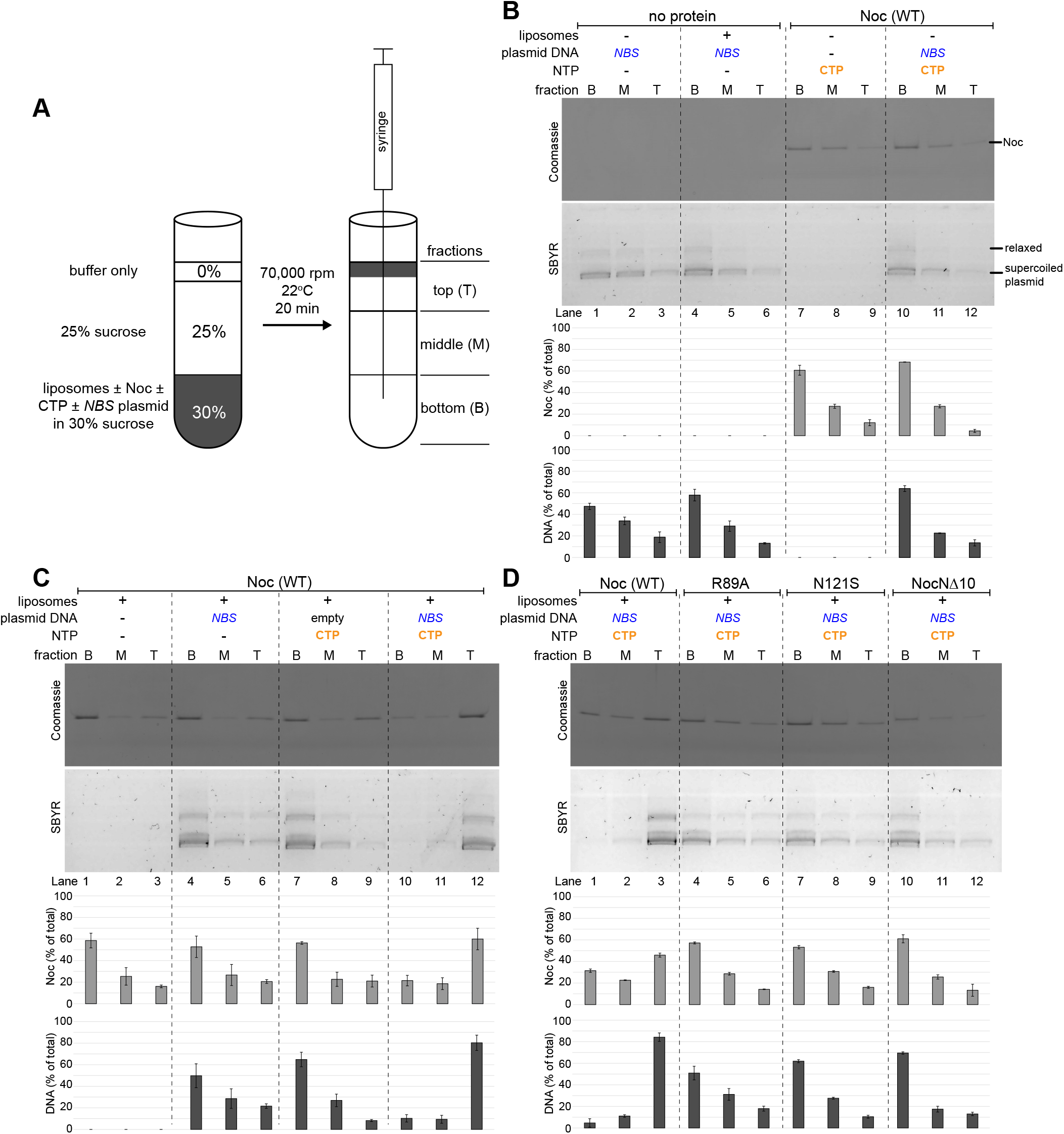
Liposome flotation assays show Noc can recruit *NBS* plasmid to the membrane in the presence of CTP, related to Figure 4. **(A)** The principle of a liposome flotation assay. Liposomes ± purified Noc ± CTP ± *NBS* plasmid were incubated in a 30% sucrose binding buffer. Buffer with 25% sucrose and 0% sucrose were subsequently layered on top sequentially. After ultracentrifugation, liposomes and associated protein/DNA migrate along the sucrose gradient i.e. floating to the uppermost fractions. Three equal fractions (bottom, middle, and top) were drawn out sequentially using a Hamilton syringe, and their protein and DNA contents were analyzed. **(B)** Control experiments: liposome flotation assays in which one component, either Noc protein, a 5-kb plasmid DNA, CTP, or liposomes, was omitted. **(C)** Analysis of *B. subtilis* Noc binding to membranes and the recruitment of plasmid DNA to membranes by a liposome flotation assay. A premix of 0.75 μM *B. subtilis* Noc ± 100 nM 5-kb plasmid DNA ± 1.0 mM CTP ± 1.0 mg/mL liposomes was incubated at 22°C for 5 min before ultracentrifugation. Either an empty plasmid or an *NBS*-harboring plasmid was employed in this assay. The resulting fractions (Bottom B, Middle M, and Top T) were analyzed by SDS-PAGE. Samples were also loaded onto a 1% agarose gel and was subsequently stained with Sybr Green for DNA. Quantification of Noc or DNA in each fraction is shown below each representative image. Error bars represent SEM from three replicates. **(D)** Other Noc variants, Noc (R89A), Noc (N121S), and NocNΔ10, were also analyzed in a liposome flotation assay. A premix of 0.75 μM *B. subtilis* Noc protein (WT or mutants) + 100 nM *NBS* plasmid + 1.0 mM CTP + 1.0 mg/mL liposomes was ultracentrifuged, and the resulting fractions were analyzed for protein and DNA contents.

**Figure S5.**
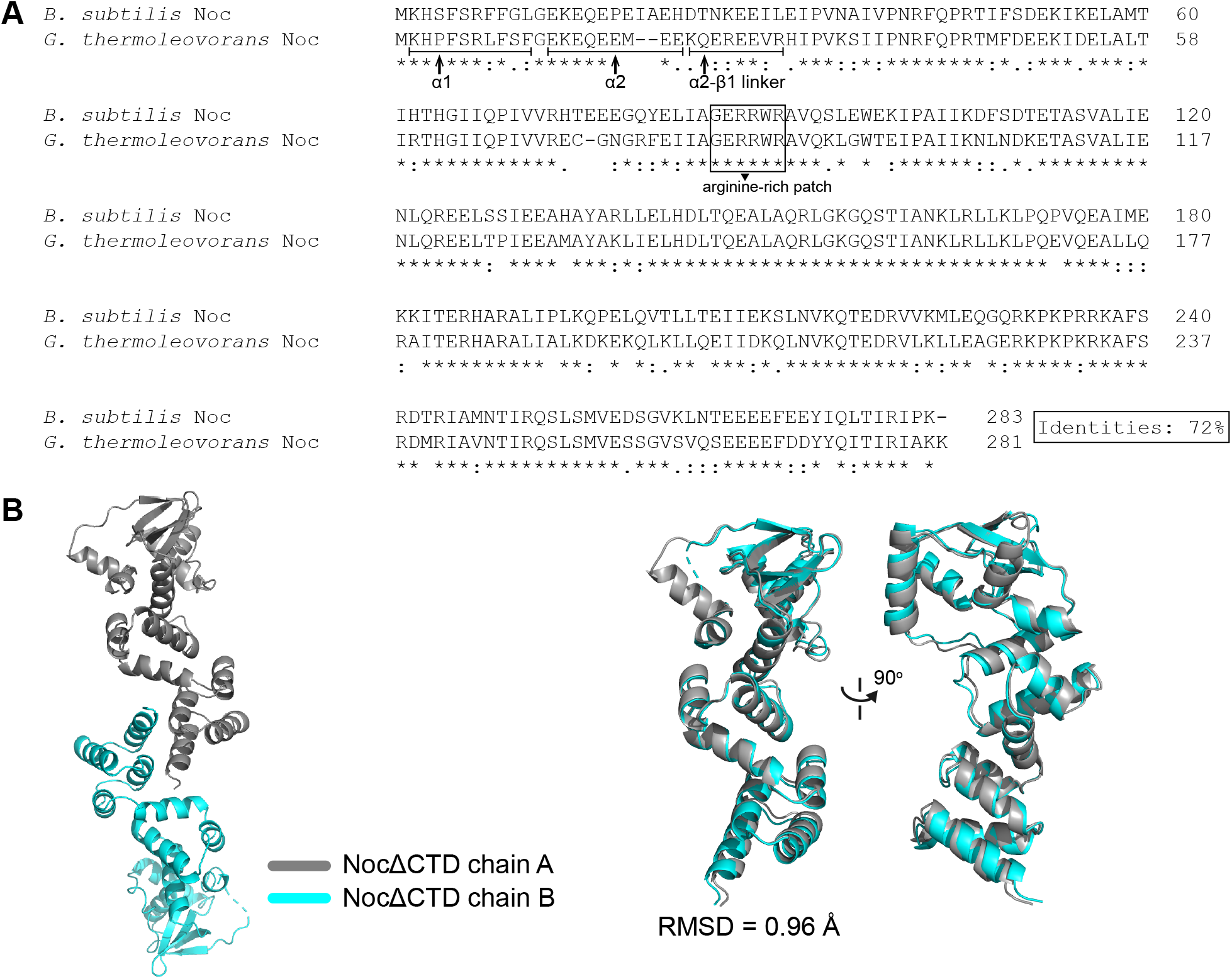
The composition of the asymmetric unit of *G. thermoleovorans* NocΔCTD crystal structure, related to Figure 5. **(A)** A sequence alignment between *B. subtilis* Noc and its homolog *G. thermoleovorans* Noc. The conserved arginine-rich patch that mediates CTP binding in *B. subtilis* ParB (Soh et al., 2019) is shown in the solid box. The sequences of helix α1, helix α2, and the α2-β1 connecting loop are also indicated on the sequence alignment. **(B)** The asymmetric unit of *G. thermoleovorans* NocΔCTD contains two copies of NocΔCTD monomers (left panel). Chains A and B are structurally similar, RMSD = 0.96 Å (right panel).

**Figure S6.**
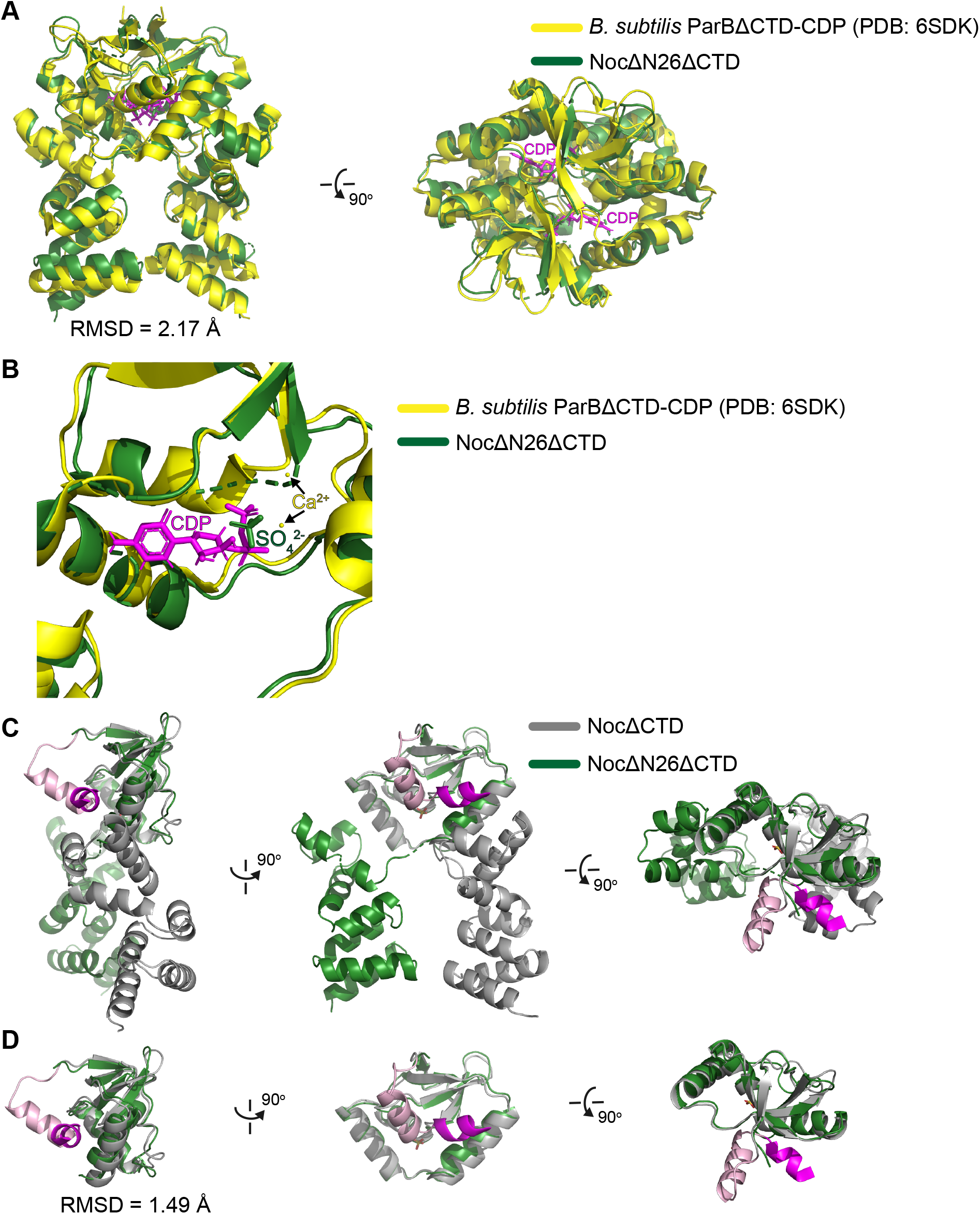
The conformation of *G. thermoleovorans* NocNΔ26ΔCTD is similar to that of a nucleotide-bound *B. subtilis* ParBΔCTD, related to Figure 6. **(A)** Superimposition between a *G. thermoleovorans* NocNΔ26ΔCTD dimer (green) and a *B. subtilis* ParBΔCTD-CDP dimer (yellow, PDB: 6SDK). CDP molecules are shown in magenta. **(B)** Magnification of the nucleotide-binding pocket of *B. subtilis* ParBΔCTD and *G. thermoleovorans* NocNΔ26ΔCTD. CDP and Ca^2+^ cations that belong to *B. subtilis* ParBΔCTD-CDP co-crystal structure are highlighted in magenta and yellow, respectively. In the *G. thermoleovorans* NocNΔ26ΔCTD structure, a sulfate ion (dark green) occupies a similar position to the β-phosphate group of CDP. **(C)** A superimposition at the N-terminal domains of a NocΔCTD monomer (grey) and a NocNΔ26ΔCTD monomer (green). The amphipathic helix α1 and helix α2 are shown in magenta and pink, respectively. **(D)** Same as panel B, but only the N-terminal domain (NTD) is shown for clarity.

**Figure S7.**
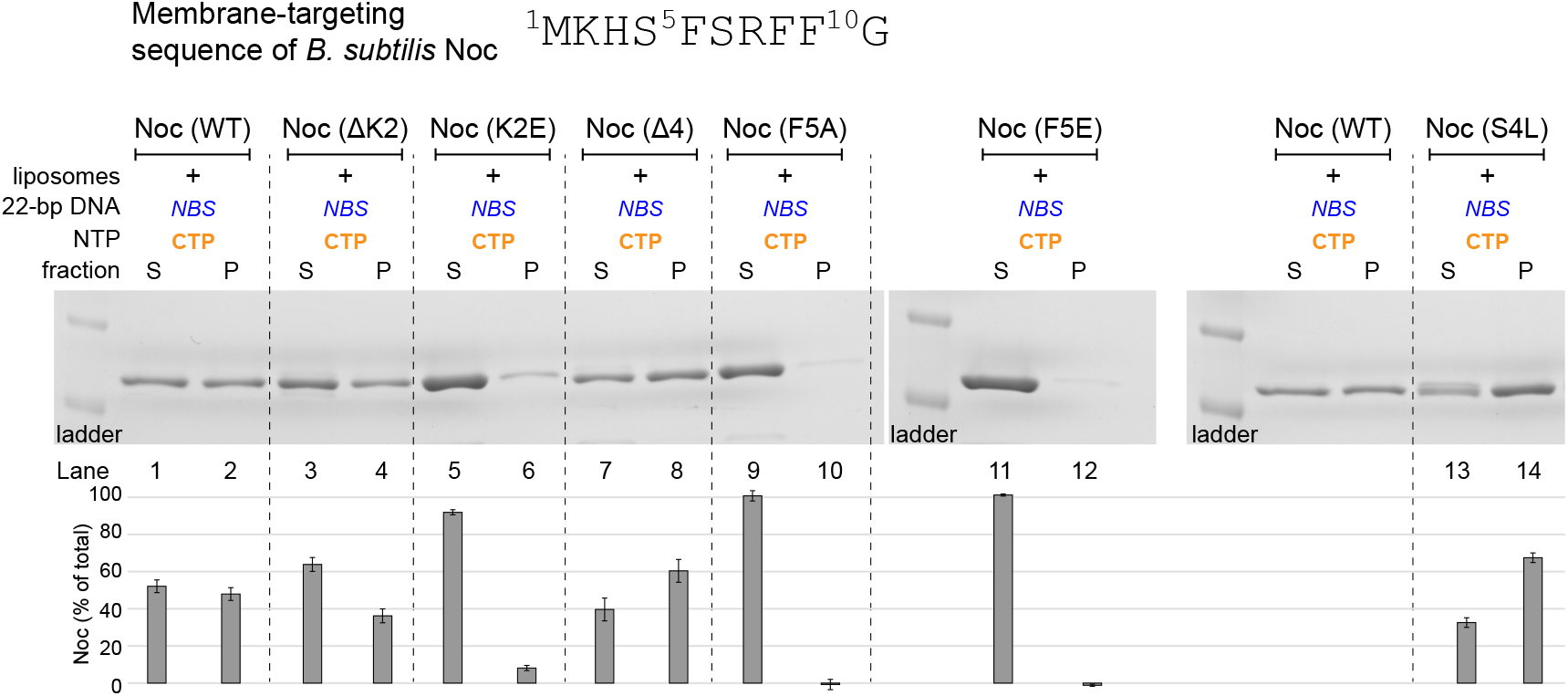
Effects of N-terminal substitutions and deletions on *B. subtilis* Noc-liposomes interaction, related to Figure 6. A premix of 1.0 μM 22-bp linear *NBS* DNA + 1.0 μM *B. subtilis* Noc protein (WT/mutants) + 1.0 mM CTP + 1.0 mg/mL liposomes was incubated at 22°C before ultracentrifugation. The resulting supernatant (S) and pellet (P) fractions were analyzed by SDS-PAGE. Quantification of Noc in each fraction is shown below each representative image. Error bars represent SEM from three replicates.

**ABLE S1.** Bacterial strains.

| Strains | Description | Source |
| --- | --- | --- |
| *E. coli* Rosetta (DE3) | F<sup>-</sup> *ompT hsdS<sub>B</sub>(r<sub>B</sub><sup>-</sup> m<sub>B</sub><sup>-</sup>) gal dcm* (DE3) pRARE (chloramphenicol<sup>R</sup>) | Merck |
| DWA117 | *B. subtilis* 168CA (*trpC2*) Δ*noc::tet* | (Adams et al., 2015) |
| DWA206 | *B. subtilis* 168CA (*trpC2*) Δ*noc::tet* Ω*amyE::[spc Pxyl-noc (WT)-myfp]* | (Adams et al., 2015) |
| DWA382 | *B. subtilis* 168CA (*trpC2*) Δ*noc::tet* Ω*amyE::[spc Pxyl-nocNΔ10-myfp]* | (Adams et al., 2015) |
| DWA546 | *B. subtilis* 168CA (*trpC2*) Δ*noc::tet* Ω*amyE::[spc Pxyl-noc (R89A)-myfp]* | (Adams et al., 2015) |
| 4746 | *B. subtilis* 168CA (*trpC2*) Δ*noc::tet* Ω*amyE::[spc Pxyl-noc (N121S)-yfpmut1]* | This study |
| DWA564 | *B. subtilis* 168CA (*trpC2*) Δ*noc::tet* Δ*minCD::kan* Ω*amyE::[spc Pxyl-noc (WT)-myfp]* | (Adams et al., 2015) |
| DWA566 | *B. subtilis* 168CA (*trpC2*) Δ*noc::tet* Δ*minCD::kan* Ω*amyE::[spc Pxyl-nocNΔ10-myfp]* | (Adams et al., 2015) |
| DWA600 | *B. subtilis* 168CA (*trpC2*) Δ*noc::tet* Δ*minCD::kan* Ω*amyE::[spc Pxyl-noc (R89A)-myfp]* | (Adams et al., 2015) |
| 4747 | *B. subtilis* 168CA (*trpC2*) Δ*noc::tet* Δ*minCD::kan* Ω*amyE::[spc Pxyl-noc (N121S)-yfpmut1]* | This study |

**TABLE S2.**
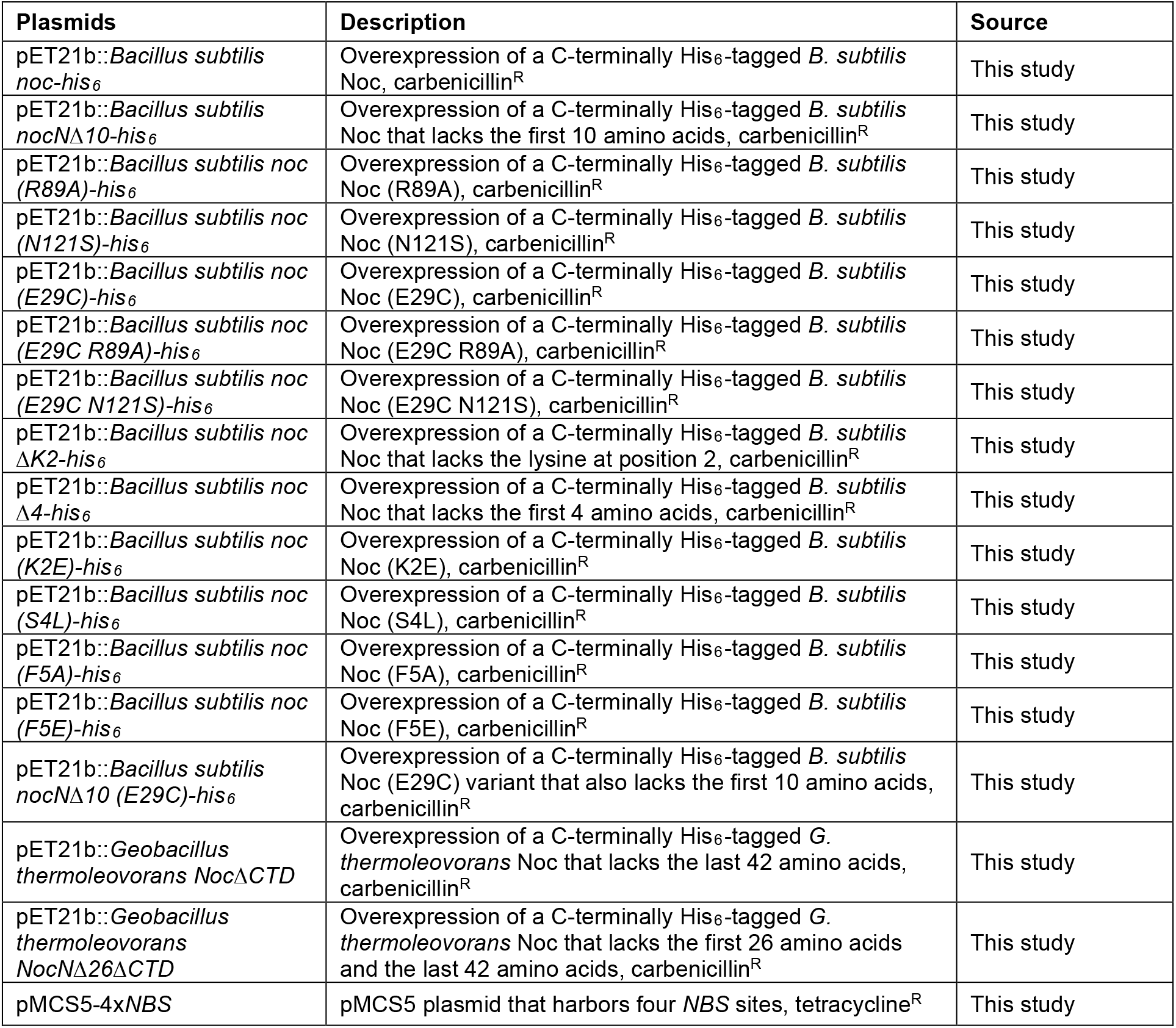
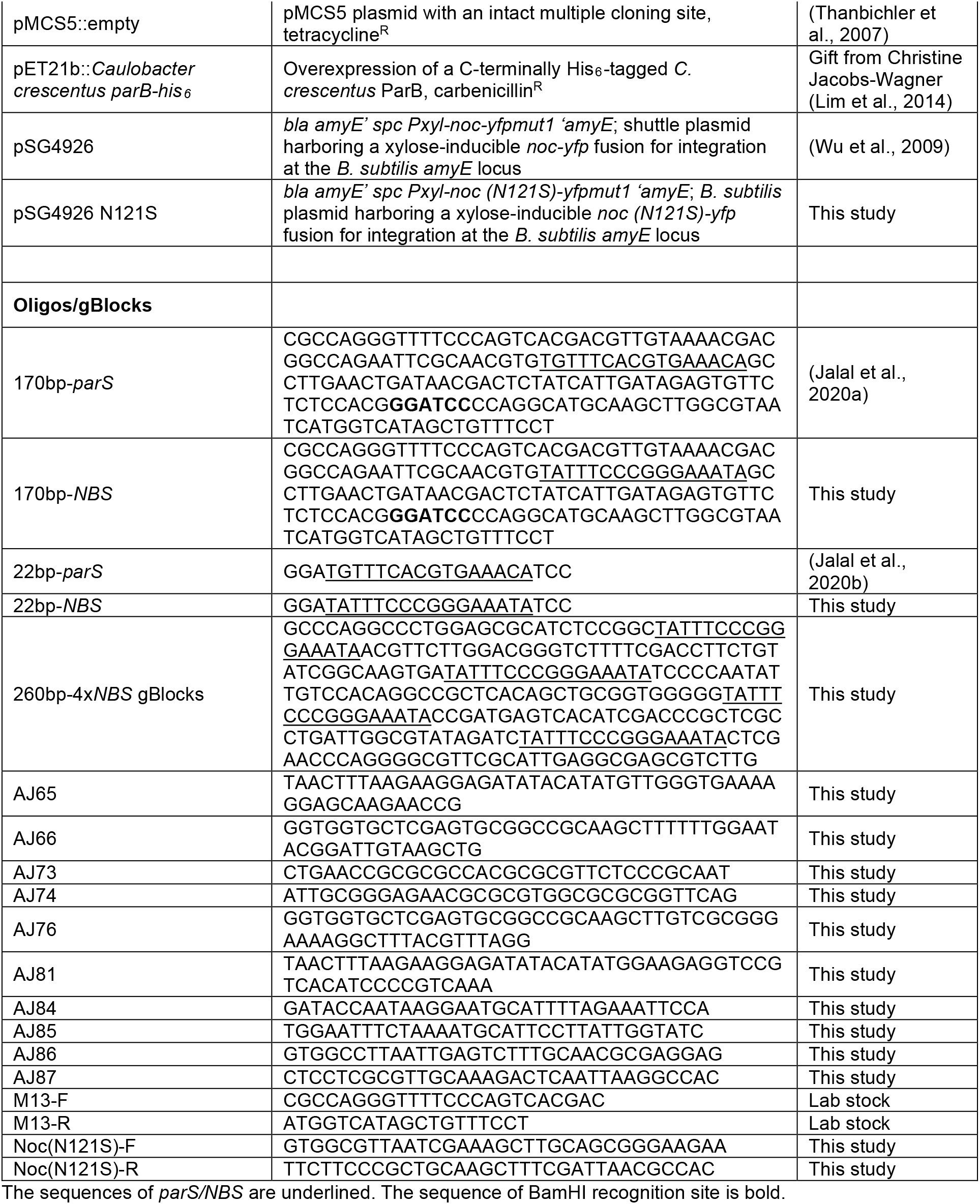
DNA, plasmids, and oligos.

**TABLE S3.** X-ray data collection and processing statistics.

| Structure | *G. thermoleovorans*<br>NocΔCTD - iodide | *G. thermoleovorans*<br>NocΔCTD - native | *G. thermoleovorans*<br>NocNΔ26ΔCTD |
| --- | --- | --- | --- |
| *Data collection* |  |  |  |
| Diamond Light Source beamline | I04 | I04-1 | I04 |
| Wavelength (Å) | 1.800 | 0.912 | 0.980 |
| Detector | Eiger2 XE 16M | Pilatus 6M-F | Eiger2 XE 16M |
| Resolution range (Å) | 96.61 – 3.40 (3.67 – 3.40) | 84.89 – 2.50 (2.60 – 2.50) | 37.44 – 2.95 (3.13 – 2.95) |
| Space Group | *P*2<sub>1</sub>3 | *P*2<sub>1</sub>3 | *C*222<sub>1</sub> |
| Cell parameters (Å/°) | *a* = *b* = *c* = 136.6 | *a* = *b* = *c* = 146.8 | *a* = 105.1, *b* = 106.6, *c* = 42.2 |
| Total no. of measured intensities | 2147484 (426036) | 1458851 (163435) | 66966 (10641) |
| Unique reflections | 11990 (2445) | 36704 (4119) | 5285 (835) |
| Multiplicity | 179.1 (174.2) | 39.7 (39.7) | 12.6 (12.0) |
| Mean *I*/$\sigma$(*I*) | 15.9 (3.3) | 28.5 (1.8) | 5.5 (1.5) |
| Completeness (%) | 100.0 (100.0) | 100.0 (100.0) | 100.0 (100.0) |
| *R*<sub>merge</sub><sup>a</sup> | 0.500 (3.392) | 0.091 (2.654) | 0.281 (1.343) |
| *R*<sub>meas</sub><sup>b</sup> | 0.501 (3.402) | 0.093 (2.688) | 0.293 (1.399) |
| *CC*<sub>½</sub><sup>c</sup> | 0.999 (0.894) | 1.000 (0.670) | 0.997 (0.885) |
| Wilson *B* value (Å<sup>2</sup>) | 88.7 | 68.0 | 38.2 |
| *Refinement* |  |  |  |
| Resolution range (Å) | - | 84.89 – 2.50 (2.57 – 2.50) | 37.44 – 2.95 (3.03 – 2.95) |
| Reflections: working/free<sup>d</sup> | - | 34843/1794 | 4752/522 |
| *R*<sub>work</sub><sup>e</sup> | - | 0.210 (0.328) | 0.267 (0.438) |
| *R*<sub>free</sub><sup>e</sup> | - | 0.240 (0.386) | 0.288 (0.443) |
| Ramachandran plot: favored/allowed/disallowed<sup>f</sup> (%) | - | 98.1/1.9/0.0 | 98.0/2.0/0.0 |
| R.m.s. bond distance deviation (Å) | - | 0.010 | 0.007 |
| R.m.s. bond angle deviation (°) | - | 1.55 | 1.33 |
| Mean *B* factors: protein/sulfate/water/overall (Å<sup>2</sup>) | - | 85/118/71/86 | 74/70/0/74 |
| **PDB accession code** |  | **7NFU** | **7NG0** |

Values in parentheses are for the outer resolution shell.
<sup>a</sup> $R_{\text{merge}} = \sum_{hkl} \sum_i |I_i(hkl) - \langle I(hkl) \rangle| / \sum_{hkl} \sum_i I_i(hkl)$ .
<sup>b</sup> $R_{\text{meas}} = \sum_{hkl} [N(N-1)]^{1/2} \times \sum_i |I_i(hkl) - \langle I(hkl) \rangle| / \sum_{hkl} \sum_i I_i(hkl)$ , where $I_i(hkl)$ is the *i*th observation of reflection *hkl*, $\langle I(hkl) \rangle$ is the weighted average intensity for all observations *i* of reflection *hkl* and *N* is the number of observations of reflection *hkl*.
<sup>c</sup> *CC*<sub>½</sub> is the correlation coefficient between symmetry equivalent intensities from random halves of the dataset.
<sup>d</sup> The dataset was split into "working" and "free" sets consisting of 95 and 5% of the data respectively. The free set was not used for refinement.
<sup>e</sup> The R-factors *R*<sub>work</sub> and *R*<sub>free</sub> are calculated as follows: $R = \sum (|F_{\text{obs}} - F_{\text{calc}}|) / \sum |F_{\text{obs}}|$ , where *F*<sub>obs</sub> and *F*<sub>calc</sub> are the observed and calculated structure factor amplitudes, respectively.
<sup>f</sup> As calculated using MolProbity (Williams et al., 2018)

## Notes

### Competing Interest Statement

The authors have declared no competing interest.

